# MetaOmGraph: a workbench for interactive exploratory data analysis of large expression datasets

**DOI:** 10.1101/698969

**Authors:** Urminder Singh, Manhoi Hur, Karin Dorman, Eve Wurtele

## Abstract

The diverse and growing omics data in public domains provide researchers with a tremendous opportunity to extract hidden knowledge. However, the challenge of providing domain experts with easy access to these big data has resulted in the vast majority of archived data remaining unused. Here, we present MetaOmGraph (MOG), a free, open-source, standalone software for exploratory data analysis of massive datasets by scientific researchers. Using MOG, a researcher can interactively visualize and statistically analyze the data, in the context of its metadata. Researchers can interactively hone-in on groups of experiments or genes based on attributes such as expression values, statistical results, metadata terms, and ontology annotations. MOG’s statistical tools include coexpression, differential expression, and differential correlation analysis, with permutation test-based options for significance assessments. Multithreading and indexing enable efficient data analysis on a personal computer, with no need for writing code. Data can be visualized as line charts, box plots, scatter plots, and volcano plots. A researcher can create new MOG projects from any data or analyze an existing one. An R-wrapper lets a researcher select and send smaller data subsets to R for additional analyses. A researcher can save MOG projects with a history of the exploratory progress and later reopen or share them. We illustrate MOG by case studies of large curated datasets from human cancer RNA-Seq, in which we assembled a list of novel putative biomarker genes in different tumors, and microarray and metabolomics from *A. thaliana*.

## Introduction

Public data repositories store petabytes of raw and processed data produced using microarray (1), RNA-seq (2), and mass spectrometry for small molecules (3) and proteins (4). These data represent multiple species, tissues, genotypes, and conditions; some are the results of groundbreaking research. Buried in these data are biological relationships among molecules that have never been explored. Integrative analysis of data from the multiple studies representing diverse biological conditions is key to fully exploit these vast data resources for scientific discovery (5, 6). Such analysis allows efficient reuse and recycling of these available data and its metadata (1, 5, 7, 8). Higher statistical power can be attained with bigger datasets and the wide variety of biological conditions can reveal the diverse transcription spectrum of genes. Yet, despite the availability of such vast data resources, most bioinformatic studies use only a limited amount of the available data.

A common goal of analyzing omics data is to infer functional roles of particular features by investigating differential expression and coexpression patterns. A wide variety of R-platform tools can provide specific analyses (9–16). Such tools are based upon rigorous statistical frameworks and produce accurate results when the model assumptions hold. Some tools avoid the need to code by providing “shiny” interfaces (17) to various subsets of R’s functionalities (10, 18– 20). They can only apply selected R packages to the data, and have the general limitations that they are not well suited for very large datasets, and have very limited interactivity.

Presently, there are very limited options for researchers to interact with expression datasets using the fundamental principles of exploratory data analysis (21). Exploratory data analysis is a technique to gain insight into a dataset, often using graphical methods. Exploratory data analysis can reveal complex associations, patterns or anomalies within data at different resolutions. By adding interactivity for visualizations and statistical analyses, the process of exploration is simplified by enabling researchers to directly explore the underlying, often complex and multidimensional data. Researchers in diverse domains (e.g., experts in Parkinson’s disease, malaria, or nitrogen metabolism) can mine and remine the same data, extracting information pertinent to their areas of expertise and deriving testable hypotheses. These hypotheses can inform the design of new laboratory experiments. Being able to explore and interact with the data becomes even more critical as datasets become larger. The information content inherent in the vast stores of public data is tremendous. Due to the sheer size and complexity of such big data, there is a pressing requirement for effective interactive analysis and visualization tools (22, 23).

Increasing the usability of the vast data resources by enabling efficient exploratory analysis would provide a tremendous opportunity to probe the expression of transcripts, genes, proteins, metabolites and other features across a variety of different conditions. Such exploration can generate novel hypotheses for experimentation, and hence provide a fundamental understanding of the function of genes, proteins, and their roles in complex biological networks (6, 8, 24–28).

Here, we present MetaOmGraph (MOG), a software written in Java, to interactively explore and visualize large datasets. MOG overcomes the challenges posed by the size and complexity of big datasets by efficient handling of the data files, using a combination of data indexing and buffering schemes. Further, by incorporating metadata, MOG adds extra dimensions to the analyses and provides flexibility in data exploration. Researchers can explore their own data on their local machines. At any stage of the analysis, a researcher can save her/his progress. Saved MOG projects can be shared, reused, and included in publications. MOG is user-centered software, designed for all types of data, but specialized for expression data.

## MATERIALS AND METHODS

### A. Overview

MOG is a standalone program that can run on any operating system capable of running Java (Linux, Mac and Windows). Access to MOG easy. MOG runs on the researcher’s computer and thus the researcher does not need to rely on internet accessibility, and is not slowed down by the latency of transfer of big data. Furthermore, the data in a researcher’s project is secure, remaining on the researcher’s computer, particularly important for confidential data such as human RNA-Seq. MOG’s modular structure has been carefully designed to enable developers to easily implement new statistical analyses and visualizations in future. MOG’s Graphical User Interface (GUI) is the central component through which all the functionality is accessed (Figure 1).

**Fig. 1.**
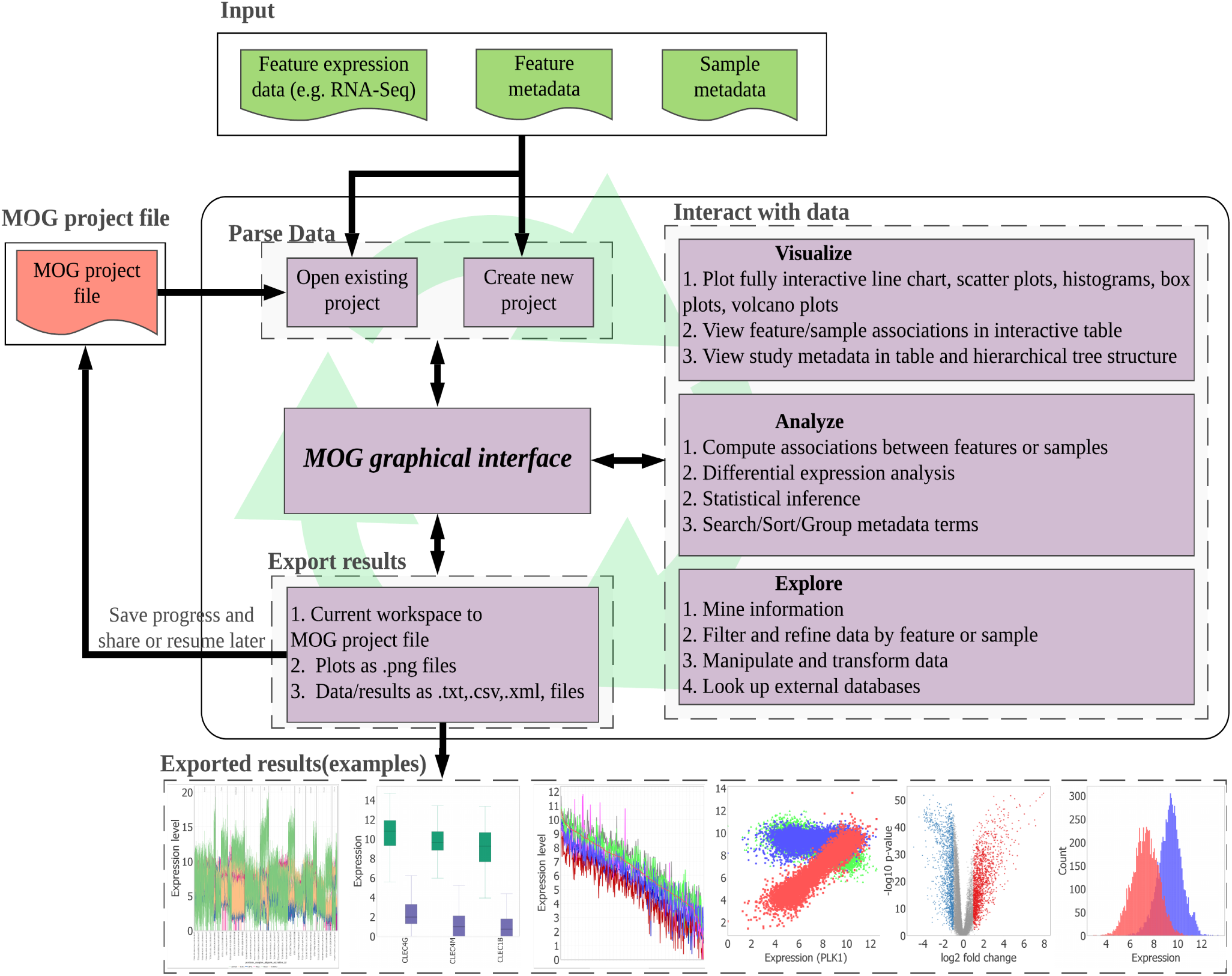
An overview of MOG’s modules. All functionality is accessed through MOG’s graphical user interface (GUI). First, the researcher selects an existing MOG project or creates a new MOG project (.mog) with input data files. Once the project is open in MOG, the workflow is non-linear. The GUI enables interactive exploration of data through various statistical analyses and visualizations. The researcher can export visualizations and results throughout the analysis, and can save and/or export her/his feature lists and statistical analyses in the MOG project file for future exploration. Saved MOG projects can be shared and further analyzed by new researchers.

#### A.1. Interactive data exploration

MOG displays all data in interactive tables and trees, providing a flexible and structured view of the data. The user can interactively filter or select data for analysis. Multiple queries can be combined and applied to discover obscure relationships within the data. This ability is particularly important for an aggregated dataset, as users may wish to split it into groups of studies, treatments, or tissues. A novel aspect of MOG is its capability of producing *interactive* visualizations. The researcher can visualize data via line charts, histograms, box plots, volcano plots, scatter plots and bar charts, each of which is programmed to allow real-time interaction with the data points and the metadata. Users can group, sort, filter, change colors and shapes, zoom, and pan interactively, via the GUI. At any point in the exploration, the researcher can search external databases: GeneCards (29), Ensembl (30), EnsemblPlants (31), RefSeq (32), TAIR (33) and ATGeneSearch (http://metnetweb.gdcb.iastate.edu/MetNet_atGeneSearch.htm) to look up additional information about the genomic features in the dataset. Researchers can also access SRA and GEO databases using the accessions present in study metadata.

#### A.2. Efficient, multithreaded and robust

A key advantage of MOG is its minimal memory usage, enabling datasets to be analyzed that are too large for other available tools. Researchers with a standard laptop or computer can easily run MOG with data files containing thousands of samples and fifty thousands of transcripts. MOG is able to perform efficiently via two complementary approaches. First, MOG indexes the data file, rather than storing the whole data in main memory. This enables MOG to work with very large files using a minimal amount of memory. Second, MOG speeds up computation using multi-threading to execute multiple tasks in parallel, optimizing the use of multi-core processors.

#### A.3. Exception handling

MOG is robust in its capability to cope with most of the many errors and exceptions (such as due to missing data or wrong data type) that can occur when handling diverse data types. If bugs are encountered, users can submit a bug report with a single click, allowing MOG developers to debug the issue.

#### A.4. Data-type agnostic

Although specifically created for the purpose of analysis of ‘omics data, which is the focus of this paper, MOG was designed to be flexible enough to generally handle numerical data and its metadata. It can interactively analyze and visualize voluminous data of any kind. Examples could be: transmission of mosquito-borne infectious diseases world-wide; public tax return data for world leaders over the past 40 years; daily sales at Dimo’s Pizza over five years; player statistics across all Women’s National Basketball Association (WNBA) teams; climate projections and outcome since 2000.

#### A.5. Leverage of third party JAVA libraries

In addition to the functionalities programmed into MOG by us, MOG borrows some existing functionality from freely available and extensively tested third-party Java libraries (JFreeChart, Apache Commons Math, Nitrite, and JDOM). This strategy results in a highly modular system that is accessible to changes and extensions.

#### A.6. Interface to R

Based on the utility and popularity of R for analysis, we have implemented an interface to facilitate execution of R scripts through MOG. By using MOG as a wrapper to R packages, the interface expands the types of analyses that can be performed. For example, a user can write an R script for hierarchical clustering of genes based on the expression levels over selected samples. The interface enables a user to interactively select/filter data using MOG and execute R scripts on selected data. This avoids writing R-code for selecting genes and samples for the analysis.

### B. Creating a new MOG project

#### B.1. Expression data files for a new MOG project

The *expression data* file contains unique identifiers (IDs) for each feature (e.g, gene) followed by information about that feature and then numerical data quantifying each feature across multiple conditions (Figure 2). For RNA-Seq analyses, each row would include a unique gene ID followed by metadata about that gene and the quantified expression level of that gene over multiple RNA-Seq runs (Figure 2). If the data is from metabolomics studies, each row would provide a unique ID for each metabolite, metadata about each metabolite, and numerical data quantifying accumulation of each metabolite over multiple GC-MS analyses. The data columns of expression data files, which contain numerical expression values, are headed by unique IDs for the corresponding samples.

**Fig. 2.**
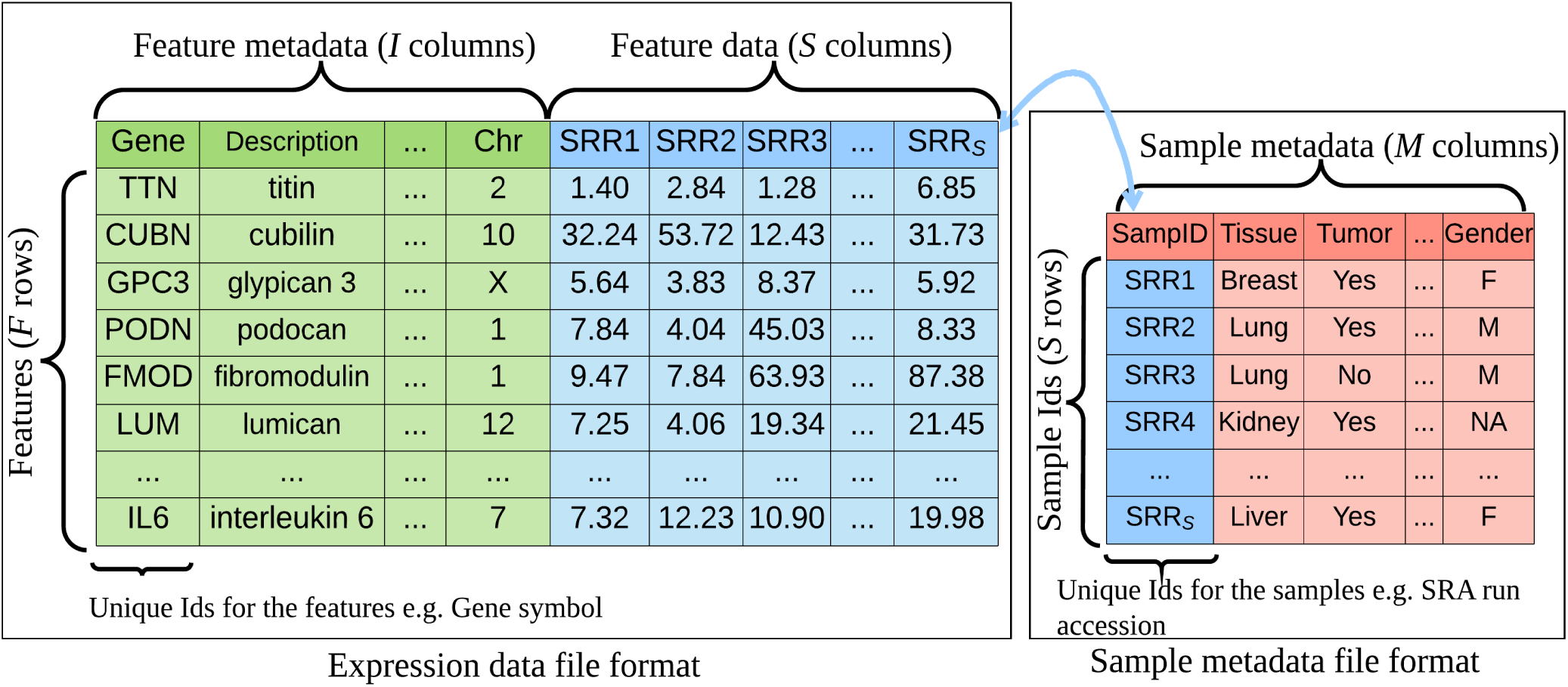
Structure of input files for a new MOG project. The *expression data file* (left table) is a matrix with *F* rows and (*I* + *S*) columns. Each row corresponds to a feature (in this example, the features are genes). The first *I* columns are feature metadata columns; the first column is a unique feature ID for each row (in this example, the gene name). The latter (*S*) columns are the expression values of *F* features over *S* samples. Each column is headed by a unique identifier for the sample. The *sample metadata file* (right table) is a matrix of *S* rows by *M* columns. Its first column contains the unique sample IDs. Thus, each row in the sample metadata file corresponds to a sample in the expression data file. The *M* columns are the metadata attributes of each analysis (e.g., each run for RNA-Seq data, each chip for microarray data).

Metadata about each feature can be added or modified by the researcher, simply by adding or modifying columns of the file (Projects provided on the MOG website can also be modified). For example, for proteomics studies, a researcher might want metadata to include: function, pathway membership, length in amino acids, phylostratum (34, 35), tertiary structure, amino acid content, and PubMed references; for metabolomics studies, metadata might include: InChi key, PubChemID, systematic name, synonyms, and molecular weight.

#### B.2. Sample metadata files for a new MOG project

This metadata is input as a delimited file (Figure 2). The first column of each row contains a unique sample ID corresponding to the sample IDs present in the data file. Other columns might contain the study ID, a description of the experimental design, experimental protocol, biological material sampled, instrument used, study abstract (Figure 2). Information can be added or modified by the user, simply by adding columns to the file. For example, a researcher might wish to add fields for references, dates, notes, or keywords.

A caveat is that leveraging archived metadata is only as good as the metadata provided; metadata about the samples may be incomplete or misleading, and the quality varies from study to study (7, 36). Despite this, metadata is key to understanding the significance of the experiments (7, 36).

The metadata for the features and samples adds extra dimensions to data exploration in MOG. For example, researchers can visualize expression of selected genes for selected treatments as compared to the other treatments (e.g., the expression of transcription factors of the NF-Y family in studies about mutants in fatty acid biosynthesis enzymes compared to mutants in enzymes of other pathways). Researchers can also filter metadata to select certain samples (e.g., samples containing the word “kidney” in the “abstract” field, and the words “stage IV” in the “tumor stage” field) for further exploration.

#### B.3. Challenges of integrating data from multiple studies

MOG is designed for exploration of complex datasets composed of multiple samples from different studies. The data supplied to MOG should be comparable across these samples. Thus, appropriate normalization should be performed on the data and effects of unwanted factors should be removed from the data (37–39). Although the steps of data normalization and integration are not a part of MOG, because the data that is input to MOG is typically from aggregate studies, we present several considerations.

Two predominate frameworks for analysis of multiple expression studies are: 1) analyze each study independently and then combine results from independent studies (meta-analysis) (5, 9, 10, 27, 40–43), or 2) combine data from multiple studies together to create a “pooled” dataset and analyze the pooled data (5, 26, 27).

In meta-analysis, if individual studies show statistically-significant results, then it is likely that the final result combined from the studies will also be significant (5). This approach avoids the major challenge of normalizing data across studies (5, 27). A caveat is that the number of samples in an individual study must be high enough to allow statistical inference (5). The major drawback of this method is that it does not allow direct comparisons across studies.

The pooling approach has the advantage that combining a large number of samples can result in better statistical power and inference; a second advantage is that data can be compared across studies (5, 44, 45). The major challenge is data normalization across studies. Despite much research on how to merge large numbers of diverse biological studies together (44, 46–49), combining heterogeneous studies, particularly of RNA-Seq data, is still a challenge because many latent unwanted factors induce variability in the data, which makes it hard to directly compare data from different samples (38, 39, 50–53).

These unwanted factors may be technical effects (e.g., library size, hardware used in sequencing, the protocol used for extraction) or unknown biological effects (e.g., a temperature drop in one of the growth rooms, or a study in which control participants were residing on a smoggy region). These unwanted factors can render the data uninterpretable in relation to its metadata, in the sense that they introduce additional covariates into the data (38, 39, 50).

If these unwanted effects are not removed, estimated associations between features after pooling data from multiple studies may be fallacious (27, 43, 54). Thus, before pooling expression data from heterogeneous studies, the effects of unwanted factors must be removed, and samples/studies that have unwieldy noise must be eliminated from the analysis.

#### B.4. Using an existing MOG project

MOG projects described in this paper and other MOG projects from well-vetted datasets are available at http://metnetweb.gdcb.iastate.edu/MetNet_MetaOmGraph.htm. Ongoing MOG project results can be saved by a researcher, regardless of whether s/he created it or it was obtained from our website. The correlations, lists, and other interactive analyses and explorations by the researcher are saved with the project. The MOG project can later be re-explored, modified, or shared.

### C. Detecting statistical association among data

Calculating statistical association between a pair of features quantifies the similarity in their expression patterns across the samples that comprise that dataset. A variety of statistical measures have been applied to estimate similarity in expression patterns (55–58). Genes with significant association may share common biological roles and pathways (26, 55, 59). Genes with significant association only under specific conditions may reveal their functional significance under those conditions (54, 60).

MOG provides the researcher with several statistical measures to estimate associations/coexpression between the features/biomolecules. It can also compute association between samples, which reflects similarity between the samples. It is up to the user to choose an appropriate method to calculate the statistical association between features or samples and interpret the results accordingly.

#### C.1. Correlation, mutual information and relatedness

We have incorporated four key methods that determine associations among features, each having its own advantages and disadvantages, depending on the types of relationships the researcher wishes to examine, and characteristics of the dataset being explored.

MOG can compute pairwise Pearson and Spearman correlation measures for selected features across samples or between selected samples across features. Pearson correlation detects the linear dependency between two random variables, *X* and *Y*, whereas, Spearman correlation coefficient measures monotonic relationships between two the variables, *X* and *Y*. Pearson correlation is computed directly using the data points; this is faster than computation of Spearman correlation, which requires converting numerical values to ranks during which some information is lost. However, Spearman correlation is less sensitive to outliers than Pearson correlation (61). Pearson or Spearman correlations values can only reveal if there is a linear or a monotonic relationship between the two random variables, which doesn’t imply causation. Pearson and Spearman correlations are often used to find coexpressed genes and generate matrices used for gene expression networks (26, 54, 56).

MOG also computes pairwise mutual information (MI) between selected features across samples. MI quantifies the amount of information shared between two random variables. MI for two discrete random variables *X* and *Y*, having the joint probability *p*(*x, y*) and marginal probabilities *p*(*x*) and *p*(*y*) respectively, is defined as:

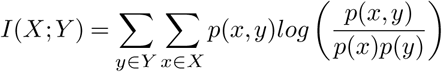

Compared to correlation methods, MI is a more general approach that can detect complex non-linear associations. The interpretation of the MI value is different than that of correlation values: an MI value of zero between two random variables implies statistical independence between them, whereas a correlation value of zero does not imply that there is no statistical dependence (61). MI has been applied to detect nonlinear associations in many gene expression datasets (28, 62– 65). MOG computes MI using B-splines density estimation, as described in Daub et.al. 2004(62).

MOG can also determine the context likelihood of relatedness (CLR) (58), estimating pairwise relatedness between features. The CLR algorithm aims to remove many false associations by comparing the MI value between a pair of features to the background distribution of MI values that include either of the features (58).

#### C.2. Meta-analysis of correlation coefficients

MOG can perform meta-analysis on Pearson correlation coefficients calculated from multiple studies. This is useful particularly when samples from different studies are not directly comparable and using pooled data for correlation is not ideal.

MOG can calculate the summary correlation coefficient from multiple correlation coefficients computed from different studies individually. Researchers can choose between a fixed effects model (FEM) or a random effects model (REM) (66, 67). The FEM combines the estimated effects by assuming that all studies probe the same correlation, i.e., studies are homogeneous, and bias is only due to study-specific variation. In contrast, the REM considers studies to be potentially heterogeneous, assuming that each study measures a biased version of the true effect and combines the estimated effects accordingly (43, 68). To perform meta-analysis of correlation coefficients with MOG, the researcher can select an appropriate column in the study metadata which identifies the different studies in the data. Then MOG computes Pearson correlation between features for samples within each study and reports the summary correlation. The researcher should carefully interpret the results, which depends on the selected model (FEM or REM) and the studies used.

### D. Differential expression between groups

Deter-mining differentially-expressed features from aggregated datasets provides direction for further data exploration. We have incorporated three statistical methods to evaluate differential expression between two independent groups of samples, and two statistical methods to compare differential expression between two paired groups of samples. For groups with independent samples, we have implemented: Mann-Whitney U test (a non-parametric test that puts no assumption about data distribution); Student’s t-test (assumes equal variance and a normal distribution of data); and Welch’s t-test (does not assume equal variance, assumes a normal distribution of data). For analysis of groups with paired samples, we have implemented in MOG: a Paired t-test (assumes normal distribution of data): and a Wilcoxon signed-rank test (a non-parametric; no assumption of data distribution).

The differential expression analyses methods implemented in MOG can achieve greater statistical power if the number of samples is sufficiently large (69). Computation and calculation of these methods via MOG is efficient and permits interactivity that promotes data exploration. For differential expression analysis of RNA-Seq/microarray data with small sample sizes, analysis in R is a viable option; methods like edger (16), DESeq2 (13) and limma (12) require raw counts as input and provide more reliable differential expression analysis for small sample sizes (70, 71). However, interactivity is limited when using these R packages.

### E. Differential correlation between groups

Features whose correlation with other features is significantly different under one environmental, genetic or developmental condition than another are differentially correlated. Differential correlation analysis is complementary to differential expression analysis and together these can reveal important features for further exploration. MOG can find the features whose Pearson correlation to a user-selected feature differs significantly in two groups of samples. To do this, MOG applies a Fisher transformation (72) and performs a hypothesis test. Differential correlations of genes reflect *shifting* biological interactions among these genes or their regulators under different conditions (60, 73), thus reflecting the context-dependency of genetic networks.

### F. Statistical inference

MOG reports p-values for all the statistical tests it performs. For correlation and MI, MOG uses a permutation test to compute the p-values. MOG speeds-up this computation using multithreading, and processing the permuted datasets in parallel (Algorithm 1). MOG also provides two popular methods to adjust the p-values for multiple comparisons: Bonferroni correction (74) and Benjamini–Hochberg (BH) correction (75).

#### Algorithm 1 Permutation test for significance

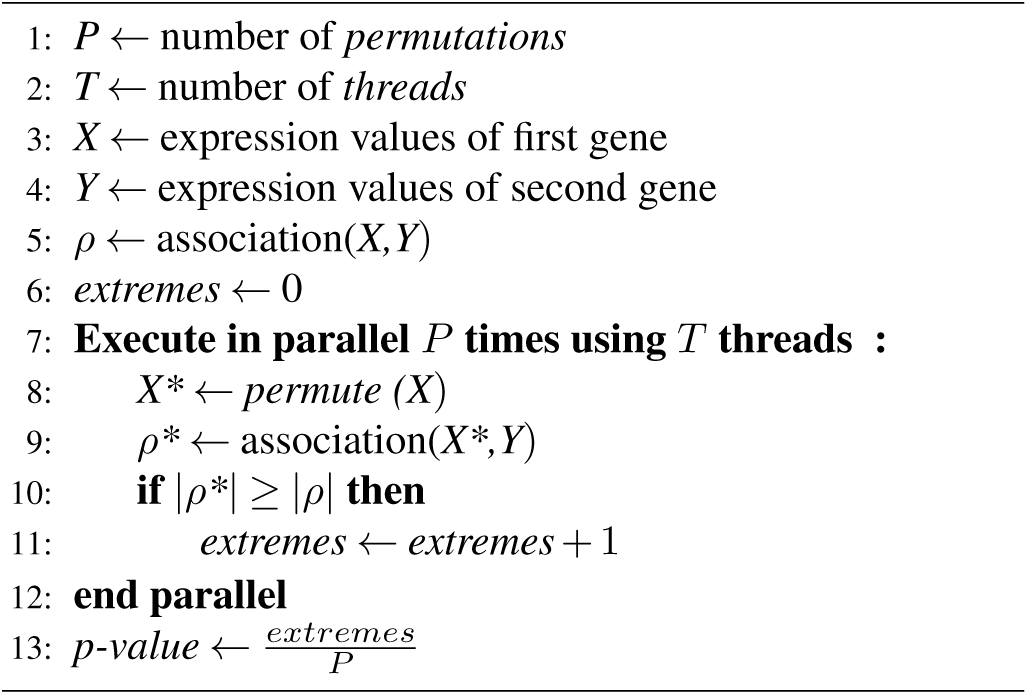

### G. Datasets

We illustrate MOG’s capabilities by exploring three diverse datasets from varied technical platforms. These are *Human cancer RNA-Seq data, A. thaliana microarray dataset* and *A. thaliana metabolomics GC-MS dataset*.

#### G.1. Human cancer RNA-seq dataset

We created a MOG project from the human cancer RNA-Seq dataset of Wang et al., 2018 (24). This highly-curated dataset combines RNA-Seq data from The Cancer Genome Atlas (TCGA, tumor and non-tumor samples) (https://cancergenome.nih.gov/) and Genotype Tissue Expression (GTEX, non-tumor samples) (76) and has been normalized and batch-corrected for study-specific biases. The resulting data is comparable across tissues (organ samples) and conditions (tumor v.s. non-tumor samples) (24) (Supplementary Table S1).

To create the MOG project, we excluded from the dataset any organ types in which the number of tumor or non-tumor samples was *<* 30. To ensure statistical independence, we included only one TCGA sample from a patient (Supplementary Table S2). We then compiled metadata for the studies/samples and for the genes in the resultant expression dataset. We downloaded the study and sample metadata (TCGA metadata from TCGAbiolinks (77); GTEX meta-data from GTEX’s website (https://gtexportal.org/home). We were unable to locate metadata for 17 of the TCGA samples and excluded these samples from the dataset (Supplementary Table S3). We extracted metadata about the genes from the HGNC (https://www.genenames.org/), NCBI Gene (https://www.ncbi.nlm.nih.gov/gene), Ensembl (30), Cancer Gene Census (78) and OMIM (79) databases, and added these information to the gene metadata in our dataset. 1,870 genes whose expression was not reported in any of the samples were removed from the dataset (Supplementary Table S4).

The final dataset used for the MOG project (Human_cancer.mog) contained expression values for 18,212 genes, 30 fields of metadata detailing each gene, across 7,142 samples representing 14 different cancer types and associated non-tumor tissues (Table 1) integrated with 23 fields of meta-data describing each study and sample. We used MOG to *log*_2_ transform the data for subsequent analyses.

**Table 1.**
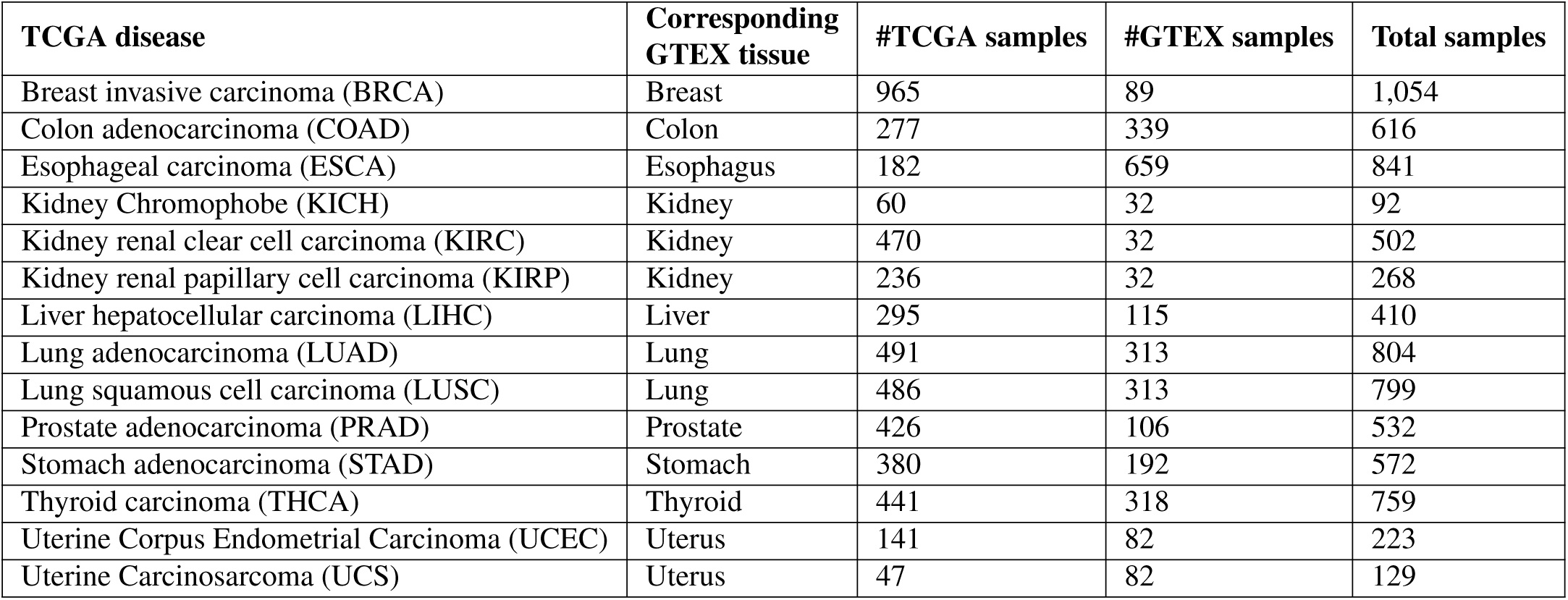
Tissue-wise summary of tumor and non-tumor samples in the human cancer RNA-Seq expression data.

#### G.2. A. thaliana microarray dataset

We created a project based on the *A. thaliana* curated microarray dataset from Mentzen and Wurtele, 2008 (26). This dataset was compiled using 963 Affymetrix ATH1 chips with 22,746 probes from 70 diverse experiments encompassing different conditions including development, stress, mutant, and other studies. All chips in the dataset were individually normalized and scaled to a common mean using MAS 5.0 algorithm. Only chips with good quality biological replicates were kept and all the biological replicates were averaged to yield 424 samples. Finally, median absolute deviation (MAD)-based normalization (80) was applied to the data. Study and sample metadata was obtained from Mentzen and Wurtele, 2008 (26). We compiled the metadata for the genes from TAIR (33) combined it with phylostrata inferences from *phylostatr* (35).

#### G.3. A. thaliana metabolomics GC-MS dataset

Small molecule composition (metabolomics) data and meta-data describing the effect of 50 mutations of genes of mostly unknown function (81) were downloaded from Plant/Eukaryotic and Microbial Resource (PMR) (82).

## RESULTS

To demonstrate MOG’s usability and flexibility, we illustrate MOG’s capabilities by exploring three diverse datasets from different perspectives. In many cases, this exploration led us to conclusions that reflect prior experimental or *in silico* results. In other cases, the exploration led us to novel predictions that could be tested experimentally. The statistical analyses and visualizations presented in this section were generated exclusively using MOG.

### H. Preliminary exploration of cancer dataset

We first used MOG to explore and assess whether the data is properly normalized and free of batch effects or not. We expect that samples from similar biological conditions should exhibit similar expression patterns for all the genes. To examine this, we computed, using MOG, pairwise Pearson correlation among samples from same conditions. All the samples had high Pearson correlation (*>* 0.70) with other samples from same biological condition, with one exception. The sample TCGA-38-4625-01A-01R-1206-07 from lung adenocarcinoma (LUAD), had unusually low expression for most genes and showed very low Pearson correlation with other LUAD samples (*<* 0.35) (Additional File 1). This sample was not included in the dataset.

Although, non-tumor samples from colon, esophagus and stomach have high Pearson correlation (*>* 0.70), we noticed the distribution of Pearson correlation values is bimodal (Supplementary Figure 1). We hypothesized that samples from these tissues were from different anatomical sites. For example, esophagus samples were from gastroesophegeal junction, muscularis and mucosa and the distribution of Pearson correlation values among samples from these three categories showed a unimodal shape (Supplementary Figure 1). Similarly, colon samples were from sigmoid colon and transverse colon and samples from sigmoid colon also showed a unimodal distribution of Pearson correlation values (Supplementary Figure 1). However, samples from transverse colon didn’t show a unimodal distribution of Pearson correlation values (Supplementary Figure 1). For stomach samples, no information about specific tissues assays was present in the metadata.

### I. Identifying a catalog of genes that are differentially expressed in 14 cancers

Using MOG, we identified differentially expressed genes, and then examined their patterns of expression across multiple samples. Initially we focused on differentially expressed genes in each cancer type with respect to the corresponding non-tumor samples. We define a gene as differentially expressed between two groups if it meets both of the following criteria:

1. Expression change is by 2-fold or more.
2. Mann–Whitney U test is significant between the two groups (BH corrected p-value *<* 10^−3^)

In each type of cancer, between 2,000-5,000 of the 18,212 genes whose expression was monitored were differentially expressed (either upregulated or downregulated) (Table 2, Supplementary Table S5-S7). Rather non-intuitively, only 16 genes were consistently upregulated and 19 genes were consistently downregulated in *every one* of the 14 cancer types. About one-third of the genes (5,748 genes) were not differentially expressed in any of the tumor types (Supplementary Table S8-S10).

**Table 2.** Number of differentiallly expressed genes found for each cancer type.

| TCGA Disease | #Upregulated | #Downregulated |
| --- | --- | --- |
| BRCA | 1,093 | 2,827 |
| COAD | 1,401 | 3,036 |
| ESCA | 1,989 | 2,229 |
| KICH | 986 | 4,214 |
| KIRC | 1,877 | 2,263 |
| KIRP | 1,152 | 2,737 |
| LIHC | 1,527 | 1,485 |
| LUAD | 1,361 | 2,753 |
| LUSC | 2,210 | 3,734 |
| PRAD | 577 | 1,633 |
| STAD | 1,527 | 1,631 |
| THCA | 993 | 1,525 |
| UCEC | 2,135 | 3,250 |
| UCS | 2,419 | 2,491 |

Several genes that are deeply implicated in cancer were not differentially expressed in any of the tumor types. One such example, tumor suppressor protein 53 (TP53) (Figure 3 A and 3 B), is thought to mediate many cancer types via mutations in its CDS that reduces its ability to suppress cancer (83); one colorectal cancer study described it as differentially expressed (fold change 1.76) (84); another did not detect differential expression (85).

**Fig. 3.**
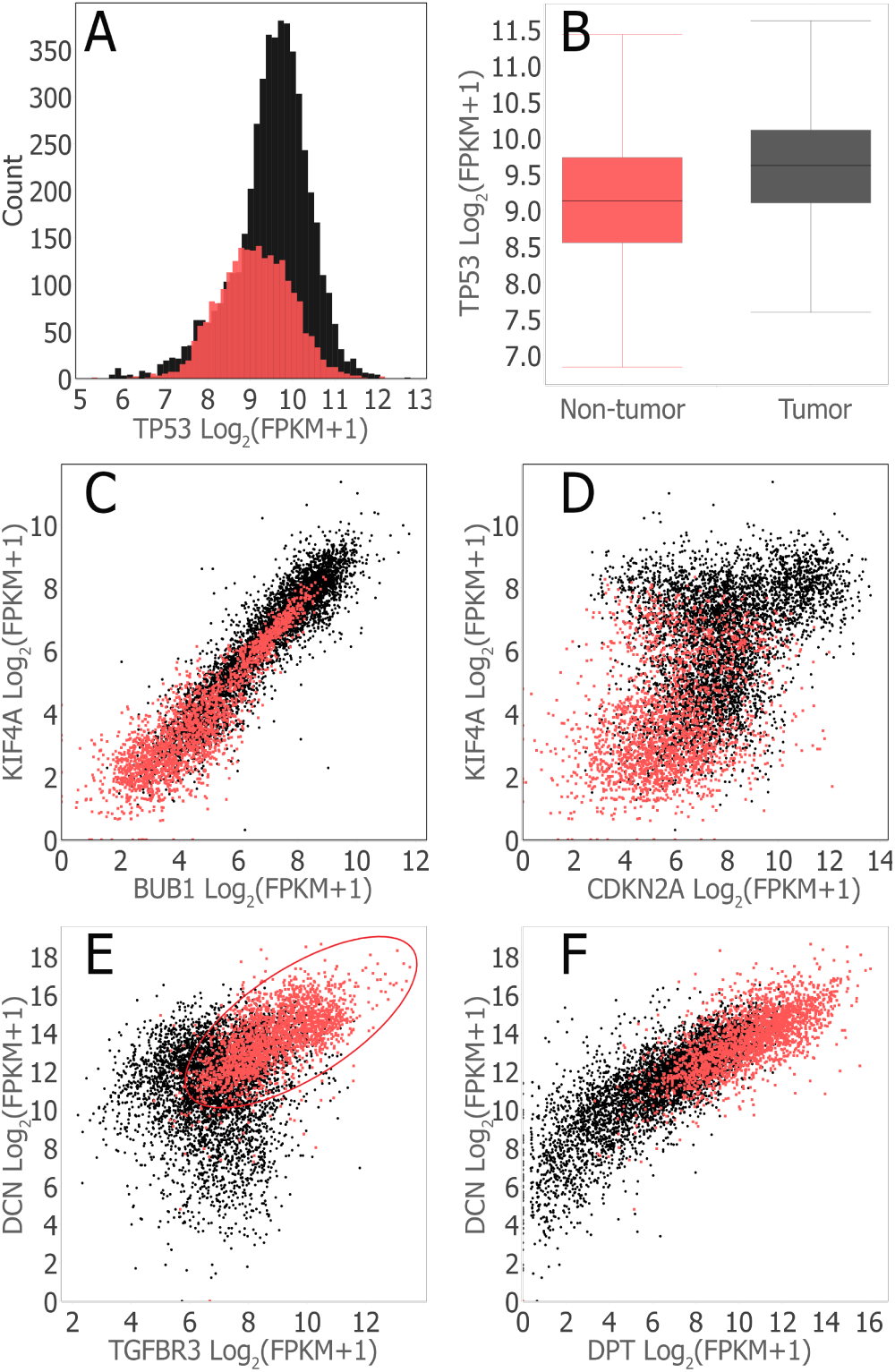
MOG visualizations of selected genes, upregulated, downregulated and un-changed across all tumor types, over all the tumor and non-tumor samples. The tumor samples are shown in black and red is used for the non-tumor samples. The plots were generated in MOG by interactively choosing the genes and the plot were split into the categories tumor and non-tumor using the sample metadata. **(A)** Histogram showing distribution of TP53 expression (number of bins parameter was set to 50). **(B)** Box plot summarizing the expression of TP53 over tumor and non-tumor samples. The horizontal line inside the box represents the median expression, which was 9.1 for non-tumor samples and 9.6 for tumor samples. **(C)** Scatter plot showing coexpression of genes BUB1 and KIF4A (both were upregulated across all tumor types). BUB1 and KIF4A show strong coexpression under both tumor and non-tumor samples (Spearman correlation = 0.92 and Spearman correlation = 0.84 respectively). Spearman correlation over both tumor and nontumor samples was 0.94. **(D)** Scatter plot showing coexpression of genes CDKN2A and KIF4A (both were upregulated across all tumor types). CDKN2A and KIF4A doesn’t show strong coexpression. The Spearman correlations were 0.36, 0.28 and 0.48 under tumor, non-tumor and all samples respectively. **(E)** Scatter plot showing coexpression pattern of genes TGFBR3 and DCN (both were downregulated across all tumor types). TGFBR3 and DCN show strong coexpression patern under non-tumor samples (Spearman correlation = 0.64). The coexpression was weak (Spearman correlation = 0.14) under the tumor samples. Spearman correlation over both tumor and non-tumor samples combined was 0.48. **(F)** Scatter plot showing coexpression pattern of genes DPT and DCN (both were downregulated across all tumor types). The Spearman correlations were 0.82, 0.69 and 0.84 under tumor, non-tumor and all samples respectively.

The expression of 15 of the 16 genes that are upregulated in all tumor types is strongly positively correlated (Spearman correlation *>* 0.60) across all tumor, non-tumor and pooled samples (tumor and non-tumor samples) (Figure 3 C, Supplementary Table S31). Cyclin dependent kinase inhibitor (CDKN2A) was the outlier (Spearman correlation *<* 0.50) (Figure 3 D; Supplementary Table S11). This may imply that these 15 upregulated genes function together closely as a module in both tumor and non-tumor cells.

In contrast, among the 19 genes that are downregulated across all cancer types, only 62 pairs are strongly correlated across all the samples (Spearman correlation *>* = 0.6) (Supplementary Table S12-S13). Of these, only 4 pairs showed strong correlation in tumor and non-tumor samples separately (Figure 3 E and 3 F; Supplementary Table S14). Seven pairs of genes are strongly correlated among tumor samples but not among the non-tumor samples (Supplementary Table S15) and 18 pairs are strongly correlated among non-tumor samples but not among the tumor samples (Supplementary Table S16), potentially reflecting a context-dependent coordination.

#### I.1. Functional analysis of differentially expressed genes

We performed gene ontology (GO) enrichment analysis using GO∷TermFinder (86) and Revigo (87) on the genes that were upregulated, downregulated or not significantly changed across all the cancer types. Upregulated genes were significantly overrepresented in GO terms related to cell proliferation: cell cycle, cell division, organelle organization, regulation of cellular component organization and regulation of cell cycle (Supplementary Table S17, Supplementary Figure 2). In contrast, downregulated genes showed no significant GO term overrepresentation; the 5,784 genes that did not change expression were enriched in GO terms: RNA processing, mRNA metabolic process, nucleic acid metabolic process, and gene expression (Supplementary Table S18, Supplementary Figure 3).

### J. Gene-level exploration

#### J.1. Expression patterns of GPC3

The Glypican 3 (GPC3) gene, encoding a glycosylphosphatidylinositol-linked heparan sulfate proteoglycan, is located on the X chromosome. GPC3 has been implicated as a critical regulator of tissue growth and morphogenesis (88). Mutations in GPC3 have been linked to Simpson-Golabi-Behmel syndrome (SGBS) and Wilm’s tumour (89–91). In embryonic development, GPC3 inhibits the cell proliferation and specialization hedgehog signaling pathway (92).

In tumors, GPC3’s role is complex and not well understood. It promotes or inhibits cell growth depending on the cancer type (93, 94). Hence, we explored expression patterns of GPC3 using MOG. Our differential expression analysis of the 14 cancer types, using Mann-Whitney test, indicates that GPC3’s expression in the LIHC samples is 30 times higher than in the non-tumor liver samples, and 8 times higher in the UCS samples compared to the non-tumor uterus samples. In contrast, GPC3 is downregulated in 9 tumor types (BRCA, COAD, ESCA, KIRC, KIRP, LUAD, LUSC, THCA, and UCEC) and unchanged in 3 tumor types (KICH, STAD and PRAD) (Figure 4 A and 4 B; Supplementary Table S19). These results are consistent with what has been reported in the literature, using different data. In liver cancer, GPC3’s expression was highly upregulated (93, 95, 96), and it has been suggested as a diagnostic biomarker and as a potential target for cancer immunotherapy in hepatocellular carcinoma (95–97). GPC3 was previously found to to be downregulated in breast (98), lung (99) and ovarian cancers (100) and its role as a tumor suppressor has been investigated in lung and renal cancer (100, 101).

**Fig. 4.**
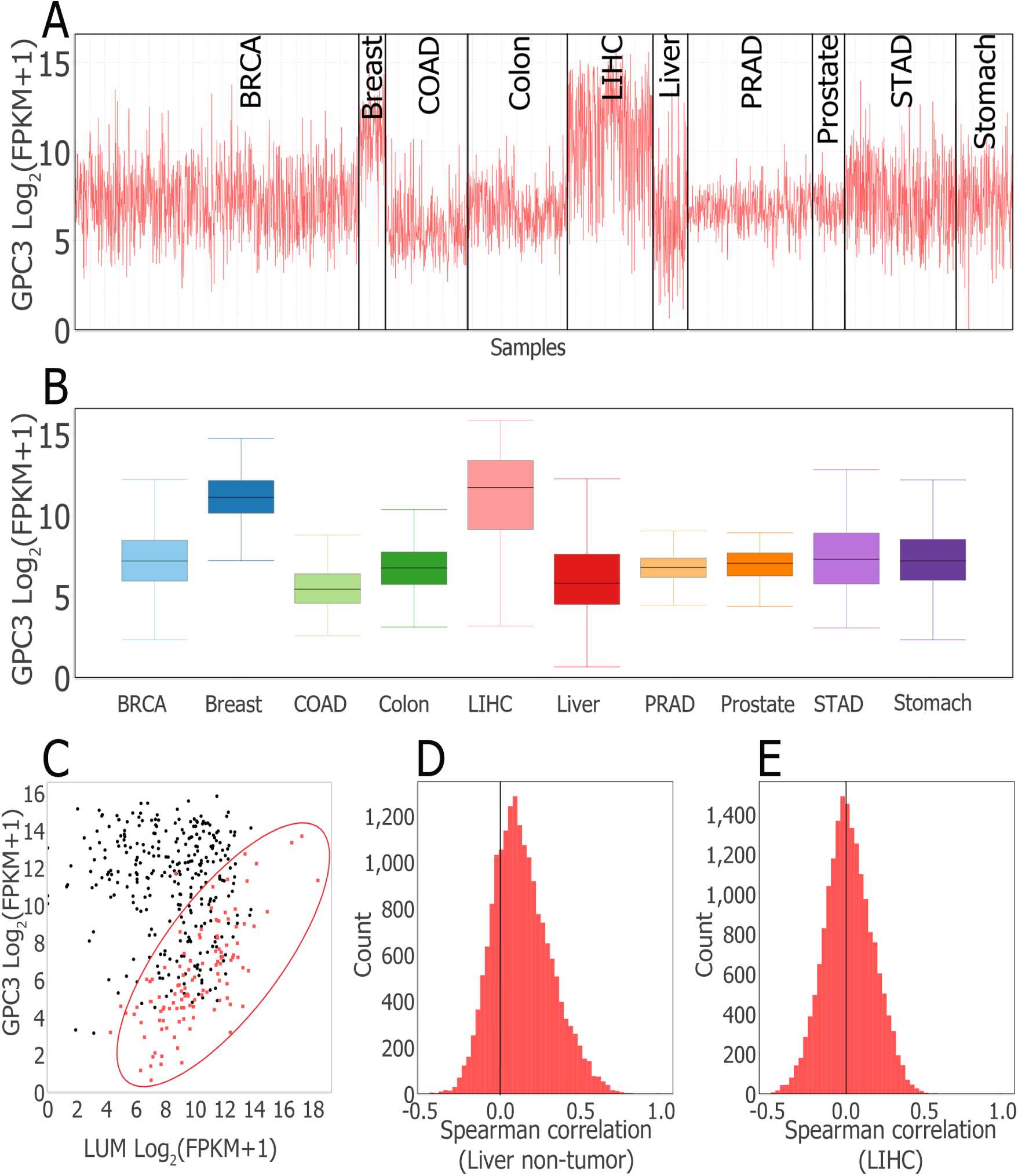
MOG visualizations of glypican 3 (GPC3) expression pattern in tumor and non-tumor tissues. **(A)** Line chart of GPC3 generated by interactively filtering by study metadata to retain 3,184 samples from 5 tumor types and corresponding non-tumor tissues and grouping the chart by tissue/tumor type. **(B)** Box plot summary of data in **(A)**. Generated by interactively splitting GPC3 box plot according to tissue/tumor type. **(C)** Scatter plot showing co-expression of GPC3 and Lumican (LUM) in liver non-tumor and LIHC samples. In non-tumor liver tissue (red), GPC3 and LUM expression are strongly correlated (Spearman correlation = 0.70). In LIHC samples (black), GPC3 and LUM expression shows no association (Spearman correlation = *−*0.10). **(D and E)** Histograms of distribution of Spearman correlation coefficients of GPC3 expression with expression of all the other genes under: non-tumor liver samples (D) and LIHC samples (E). In non-tumor liver samples the distribution of Spearman correlation coefficients has a longer right tail indicating presence of higher Spearman correlation coefficients as compared to LIHC samples. In LIHC samples the distribution of Spearman correlation coefficients is centered around zero and the correlation coefficients roughly bounded between −0.5 and 0.5.

We then investigated coexpression patterns of GPC3 in the different tumor and non-tumor tissues (Additional File 3). We selected a cutoff of Spearman correlation *>* = 0.6 or *<* = *−* 0.6 to identify genes coexpressed with GPC3. We found that GPC3’s coexpression patterns were very different across all the tumor and non-tumor samples (Figure 4 C). Genes that are correlated with GPC3 vary greatly across different tissues (Figure 4 D and 4 E; Supplementary Table S20). For example, 4,219 genes were coexpressed with GPC3 in esophagus non-tumor samples, whereas no gene was coexpressed with GPC3 in non-tumor samples from prostate and stomach (Supplementary Table S20). Coexpressed genes also differed according to whether disease was present. For 7 tissues, the number of coexpressed genes with GPC3 were lower in tumor samples than those in non-tumor samples (Supplementary Table S20). For example, although 192 genes were co-expressed with GPC3 in non-tumor liver samples, no genes were coexpressed with GPC3 in LIHC samples.

We analyzed GO term overrepresentation for GPC3 coexpressed genes from colon, esophagus, kidney and liver samples. GPC3 coexpressed genes in other non-tumor tissues were less than 10. We found that biological adhesion and cell adhesion were two GO functions present in all the sets of coexpressed genes from colon, esophagus, kidney and liver samples. The GO terms cell development, extracellular structure organization, extracellular matrix organization and multicellular organism development were present in all sets except the one from lung. Multiple other GO terms were overrepresented in a tissue specific manner (Supplementary Table S21-S26).

#### J.2. GPC3 associated modules in tumor versus non-tumor liver samples

To explore potential interactions of GPC3 with other genes in non-tumor and tumor liver samples, we used MOG to build two gene coexpression networks from the 3,012 genes that were differentially expressed in LIHC– one network from non-tumor liver samples, and a second network from LIHC samples (Supplementary File 3; Additional File 3). Then, we imported each network into Cytoscape (102) and identified the tightly connected modules using MCODE (103).

In the network built using non-tumor liver samples, GPC3 was part of a module that was ranked the second most significant by MCODE’s algorithm (73 nodes (genes); MCODE score 30.7). GPC3 was directly connected with 21 genes (Supplementary Figure 4). This module was most enriched in GO terms: sulfur compound catabolic process, glycosaminoglycan catabolic process, aminoglycan catabolic process, and extracellular matrix organization (Supplementary Table S27; Supplementary Figure 5). In contrast, GPC3 wasn’t significantly coexpressed with other genes in the LIHC samples, and thus was absent from the LIHC network. However, this network contained a module with 114 genes (MCODE score 94.3), 33 of which were in the GPC3-connected module identified from the network built using liver non-tumor samples (17 of these genes were directly connected with GPC3) (Supplementary Figure 4). GO analysis of this module indicated it was overrepresented in GO terms: extracellular matrix organization, extracellular structure organization, blood vessel development, and vasculature development (Supplementary Table S28; Supplementary Figure 6).

**Fig. 5.**
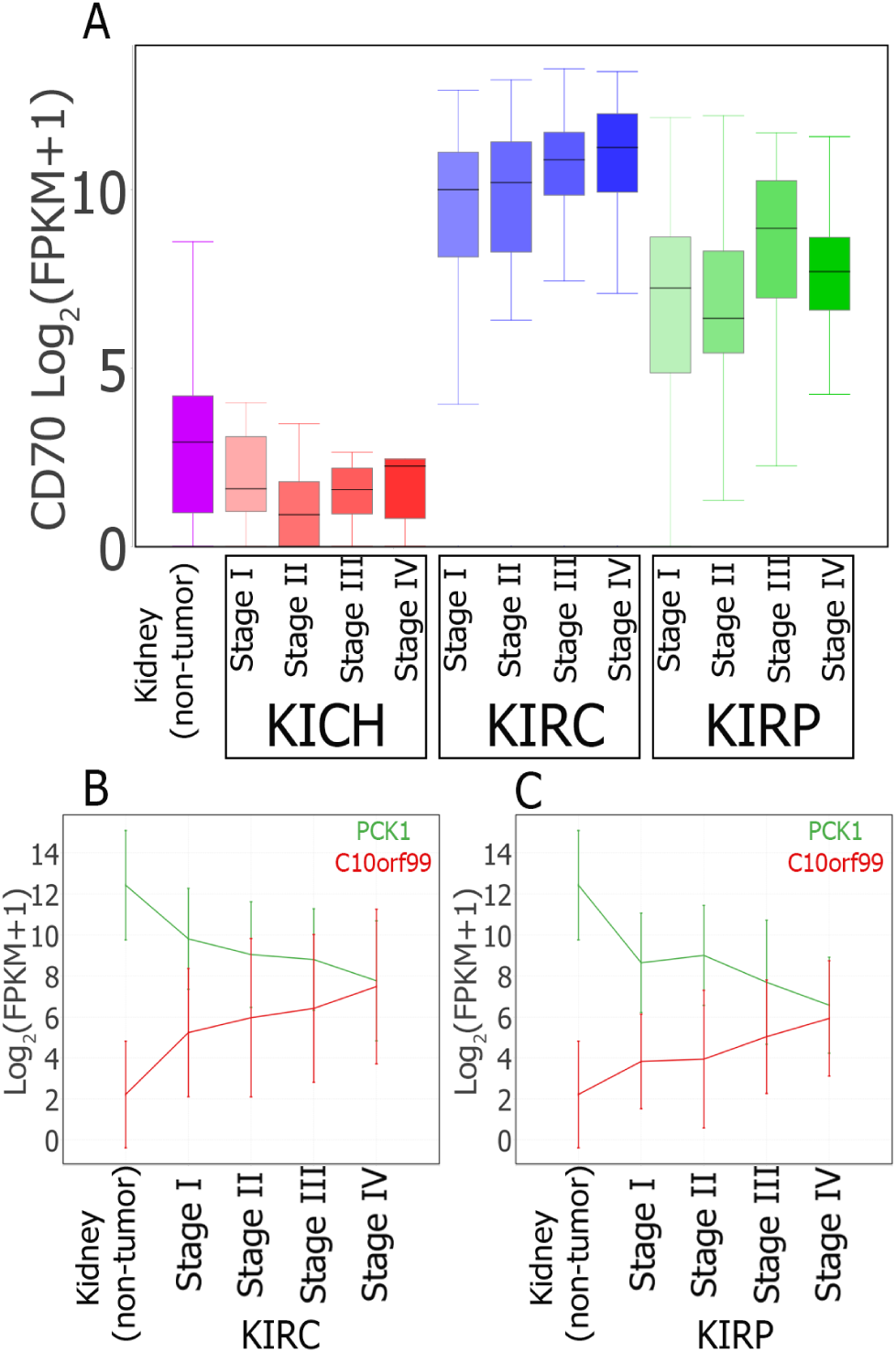
MOG visualization of expression of selected genes during progression of renal cancers. **(A)** Box plots summarizing CD70 expression in non-tumor kidney and at different stages of KICH, KIRC, and KIRP kidney cancers. CD70 is designated as prognostic unfavourable for renal cancer in THPA. CD0 levels increases 90-fold in KIRC and 14-fold in KIRP, but it is decreased in KICH (*log*_2_ *FC* = 1.56 adj p-value=0.004). **(B and C)** Line charts showing average expression of PCK1 (green) and C10orf99 (red) over different stages of KIRC (B) and KIRP (C). The vertical lines are the error bars. THPA designates PCK1 as prognostic favourable and C10orf99 as prognostic unfavourable for renal cancer.

**Fig. 6.**
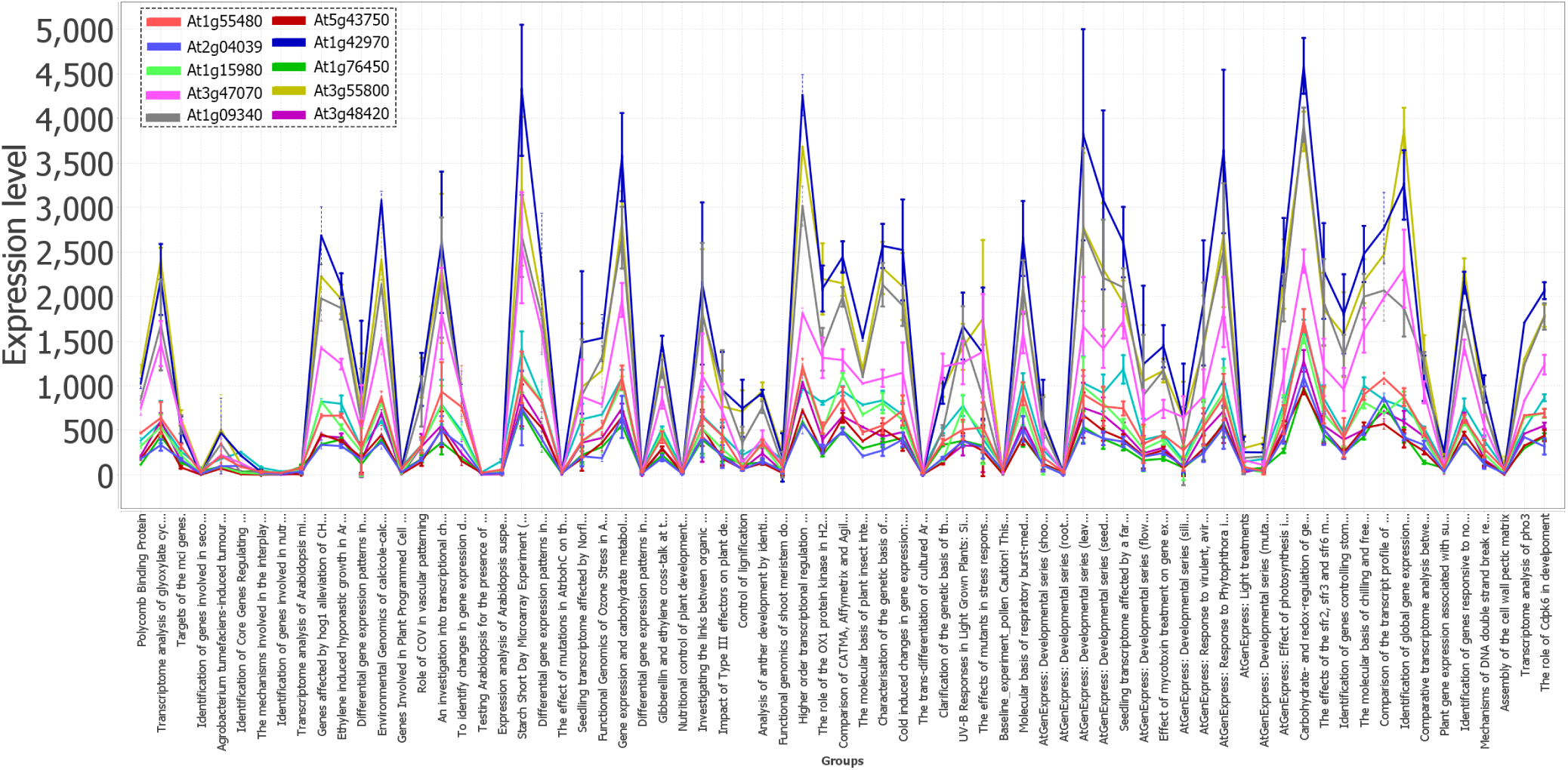
Spearman correlation followed by line-plot visualization, using MOG, shows that Met1 (At1G55480) is highly expressed in photosynthetic tissues and highly correlated with several genes of unknown function. The “peaks” of expression are all leaf samples; the “troughs” of expression are predominantly root and cell culture samples. *A. thaliana* Affymetrix microarray dataset representing 71 diverse studies and a wide variety of environmental, genetic and developmental conditions (26). Several genes of unknown function are closely coexpressed in this cluster.

### K. Stage-wise analysis of cancer data

#### K.1. Identifying new candidate biomarkers for cancers

We used MOG to identify genes whose expression is associated with disease progression; these genes are potential biomarkers. Specifically, we identified genes upregulated in the different cancer types; then, we sorted samples from each organ-type into early stage (stage I or stage II) and late stage (stage III and later) based on the study metadata. Finally we performed Mann-Whitney test and identified those that were also upregulated in late stage compared to early stage (2-fold change or more, and B-H corrected *p − value <* 0.05) from among the upregulated genes. These genes would likely show increasing expression with cancer progression. We similarly identified the genes that were downregulated in late tumor stages compared to early stage cancer. These genes would likely show a decreasing pattern of expression with cancer progression.

A total of 221 genes showed an increasing expression pattern during tumor progression (ESCA:91, KIRP:89, THCA:25, KIRC:24); 227 genes showed a decreasing expression pattern (ESCA:89, KIRP:68, LIHC:64, KIRC:13) (Supplementary Table S29; Additional File 4). We compared the results obtained by MOG to data in The Human Protein Atlas (THPA) (104). Out of the 111 genes we identified as increasing during progression of KIRC or KIRP, 56 have been described as unfavourable prognostic for renal cancer by THPA (Supplementary Table S30). For example, Figure 5 A shows expression of the Cluster of Differentiation 70 (CD70) gene in KICH, KIRC and KIRP tumors; CD70 expression increased with KIRC and KIRP progression; CD70 gene is marked by THPA as a prognostic unfavourable for renal cancer.

Out of the 79 genes we identified as decreasing with cancer progression in KIRC or KIRP, 39 were labeled as prognostic favourable for renal cancer by THPA (Supplementary Table S30). Figure 5 B and 5 C shows the expression pattern of Phosphoenolpyruvate Carboxykinase 1 (PCK1) and Chromosome 10 Open Reading Frame 99 (C10orf99) over different stages of KIRC and KIRP. Further, 27 genes out of the 64 that we identified as decreasing with cancer progression in LIHC were labeled as prognostic favourable for liver cancer in THPA (Supplementary Table S30). Out of 25 genes identified as having increasing pattern in THCA, none were labeled as prognostic by THPA (Supplementary Table S30).

We propose that the other 327 genes identified in this study not labeled as prognostic in THPA could be new potential prognostic biomarkers for these tumor types (Supplementary S30). For example, Table 3 lists genes identified by our method that are not marked as prognostic in THPA but have been experimentally studied for their potential role as a prognostic biomarker in LIHC or THCA.

**Table 3.**
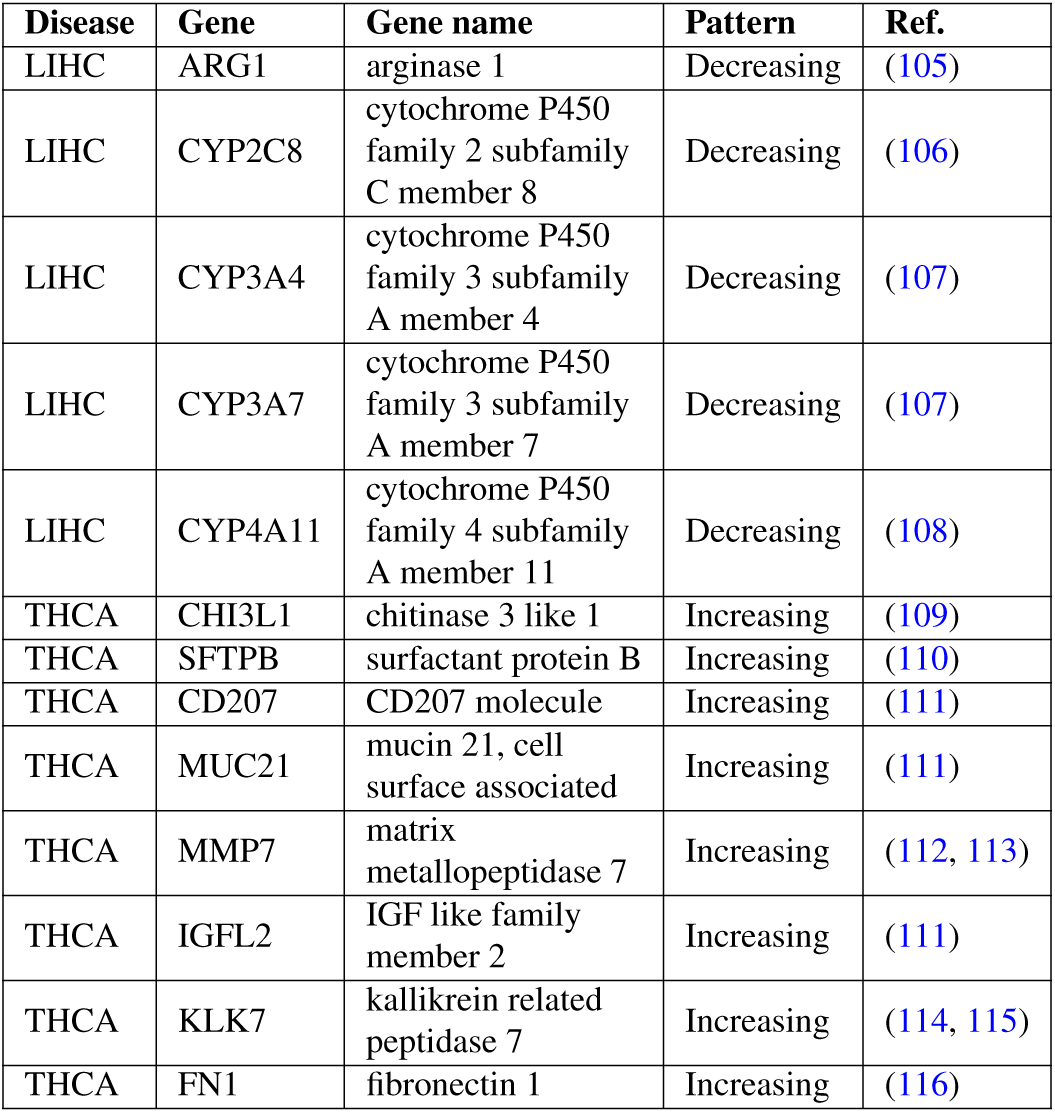
Genes identified by MOG as showing changed expression in LIHC and THCA with progression of cancer (B-H corrected *p − value <* 0.05) that have also been considered in studies (Ref. column) for use as potential prognostic biomarkers. These genes are not marked as prognostic in The Human Protein Atlas (THPA) (104) for the given cancer types.

For prognosis and personalized treatment, the *exceptions* are extremely important. By using MOG to explore very large sets of data, a researcher can visualize in more detail whether certain samples do not show changed expression of a prognostic marker; this could lead to additional experimental analysis to reveal whether something about these “no-show” samples might be biologically distinct. A researcher can also evaluate which cancers have a particular biomarker; how specific is a given biomarker to a particular type and stage of cancer. For example, ARG1, CYP2C8, CYP3A4, CYP3A7 and CYP4A11, which we identified using MOG as decreasing expression with LIHC progression, have been recently studied as prognostic markers for hepatocellular carcinoma (105–108).

### L. *Arabidopsis thaliana* microarray data

Our aim in the case study of *A. thaliana* microarray data was to explore patterns of expression of genes that little or nothing is known about. The well-vetted dataset we used (26), encompasses expression values for 22,746 genes across 423 *A. thaliana* microarray chips; the chips represent 71 diverse studies and a wide variety of environmental, genetic and developmental conditions (26). We updated the gene metadata to the current TAIR annotations (33); to these we added phylostrata designations (obtained from phylostratr (35)).

First, we sought to identify genes of unknown or partially-known function that might be involved in regulation of photosynthesis, the process that gave rise to the earths oxygenated atmosphere and the associated evolution of extant complex eukaryotic species. Assembly and disassembly of the photosystem I and II complexes are dynamic processes of photosynthesis that respond sensitively to shifts in light and other environmental factors (117–120); Met1 (AT1G55480) is a 36 Kda protein that regulates the assembly of the photosystem II (PSII) complex (120). To explore genes that might be involved in PSII assembly, we calculated Spearman correlation of expression of Met1 with that of the 22,746 genes represented on the Affymetrix chip (Figure 6). This analysis shows that expression of 104 genes is highly correlated (Spearman’s coefficient *>* 0.9) to Met1 gene expression across all conditions (Supplementary Table S31).

First, we examined whether genes of photosynthesis were over-represented among the Met1 coexpressed cohort. Among the Met1 coexpressed genes, the Gene Ontology (GO) Biological Functional terms most highly over-represented (p-value *<* 10^*−*5^) are integral to the light reactions of photosynthesis: generation of precursor metabolites and energy; photosynthetic electron transport in photosystem I (PSI); reductive pentose-phosphate cycle; response to cytokinin; and PS2 assembly (Supplementary Table S32). For example, the gene most highly correlated with Met1 is At2g04039, a gene encoding the NdhV protein, which is thought to stabilize the nicotinamide dehydrogenase (NDH) complex of PS1 (121); phylostratal analysis (35) indicates that NdhV has homologs across photosynthetic organisms, streptophyta (land plants and most green algae). Eighteen of the Met1 coexpressed genes are designated as “unknown function” or “uncharacterized’; six are restricted to Viridiplantae. These genes would be good candidates to evaluate experimentally for a possible function in photosynthetic light reaction.

Our next aim was to use MOG to directly explore an orphan gene (a gene encoding a protein unrecognizable by homology to those of other species) (34, 35, 122–124), and to determine potential processes that it might be involved in. First, we filtered each gene’s target description to retain “unknown”. From these, we filtered to retain only the phylostratigraphic designation “*Arabidopsis thaliana*”. From this gene list, we identified genes that had an expression value greater than 100 in at least five samples. We selected the orphan gene of unknown function, At2G04675, for exploratory analysis. At2G04675 encodes a predicted protein of 67 aa with no known sequence domains (domains searched using CDD (125)). A Pearson correlation analysis of the expression pattern of At2G04675 with the other genes represented on the Affymetrix chip showed 48 genes had a Pearson correlation of higher than 0.95 (Supplementary Table S31); these genes are expressed almost exclusively in pollen (the male game-tophyes of flowering plants) (Figure 7). The exploration implicates At2G04675 as a candidate for involvement in some aspect of pollen biology.

**Fig. 7.**
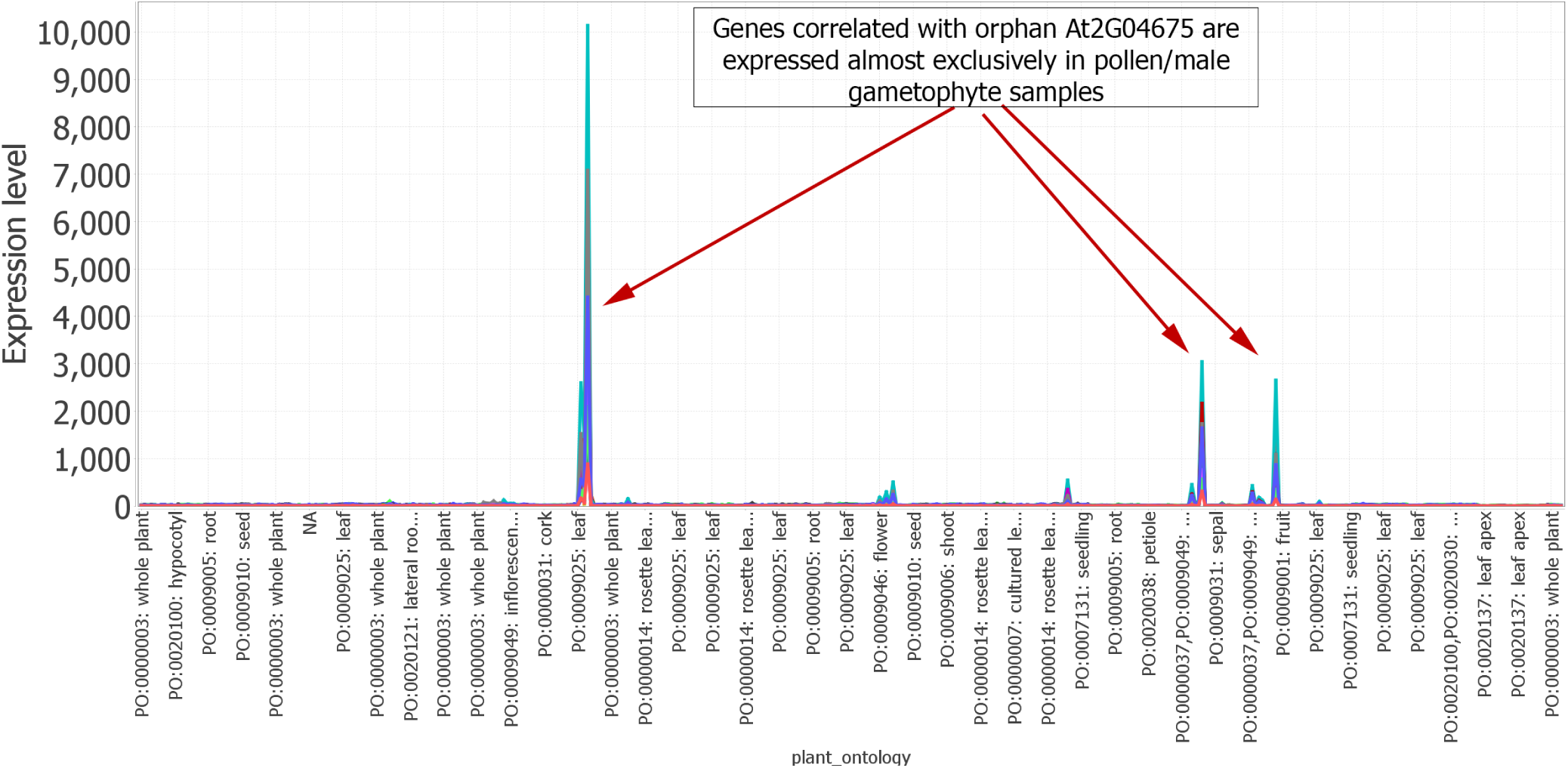
MOG line chart visualization shows the orphan gene At2G04675, of no known function, and genes highly correlated with At2G04675 are expressed almost exclusively in pollen/male gametophyte samples. *A. thaliana* Affymetrix microarray dataset representing 71 diverse studies and a wide variety of environmental, genetic and developmental conditions (26). X-axis are samples, and Y-axis indicates their expression value. Each line represents a gene. (Lines in this visualization are for clarity and the connections from sample to sample do not imply a relationship)

Using MOG to further explore genes that are associated with pollen, we identified sets of leaf and pollen samples (Supplementary Figure 7; Additional File 5), and then calculated genes that are differentially expressed in the leaf samples versus the pollen samples using a Mann-Whitney test (Expression change is by 2-fold or more; B-H corrected p-value *<* 10^*−*3^) (Additional File 5). The GO terms most highly enriched (p-value *<* 10^*−*20^) among genes upregulated in pollen are processes of cell cycle, mitosis, organellar fission, chromosome organization and DNA repair (Additional File 5). This reflects and emphasizes the critical role of these processes in the development of male gametophyte development, particularly sperm biogenesis. Each angiosperm pollen grain must produce two viable sperm each used in the double fertilization of the ovule. Above all else, proper mitogenesis is essential to the function of a pollen grain. We visualized the *leaf vs pollen* differential analysis by volcano plot (Figure 8; Supplementary Figure 8), this time to explore genes upregulated in leaf. Among these is At1G67860, an Arabidopsis specific gene encoding a protein of unknown function. We used MOG to correlate expression of this gene versus all genes across all samples. One hundred sixteen genes, dispersed across all five chromosomes, are coexpressed with At1G67860 (Spearman correlation 0.65-0.95) (Supplementary Table S31). The genes are expressed almost exclusively in mature leaf (Supplementary Figure 9). Most have no known function; a GO enrichment test indicates that GO biological processes overrepresented among the genes are: defense response, response to stress, response to external biotic stimulus and response to other organism (p-value *<* 1.E-03) (Supplementary Table S33).

**Fig. 8.**
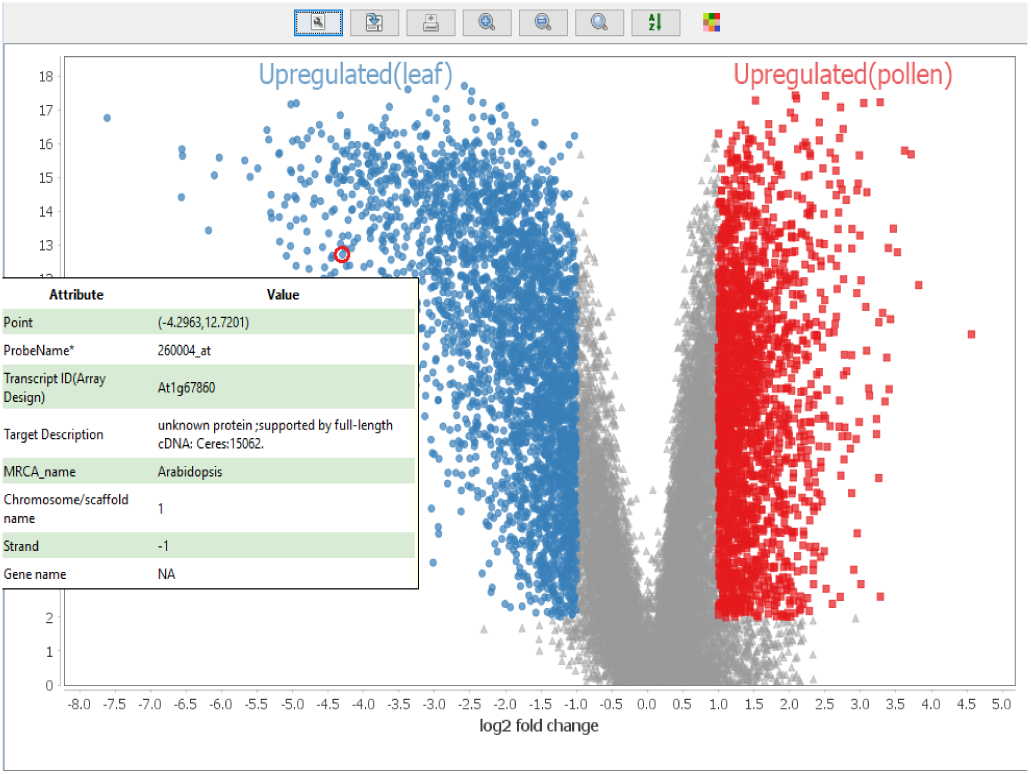
Using MOG for differential expression analysis of leaf and pollen samples, followed by volcano plot visualization. Gene metadata, revealed upon hovering the mouse over a data point, shows At1G67860, an Arabidopsis specific gene with no known function, is 16-fold more highly accumulated in leaves relative to pollen (Mann-Whitney U test; B-H corrected p-value*<* 10^*−*3^). Y axis, p-value

**Fig. 9.**
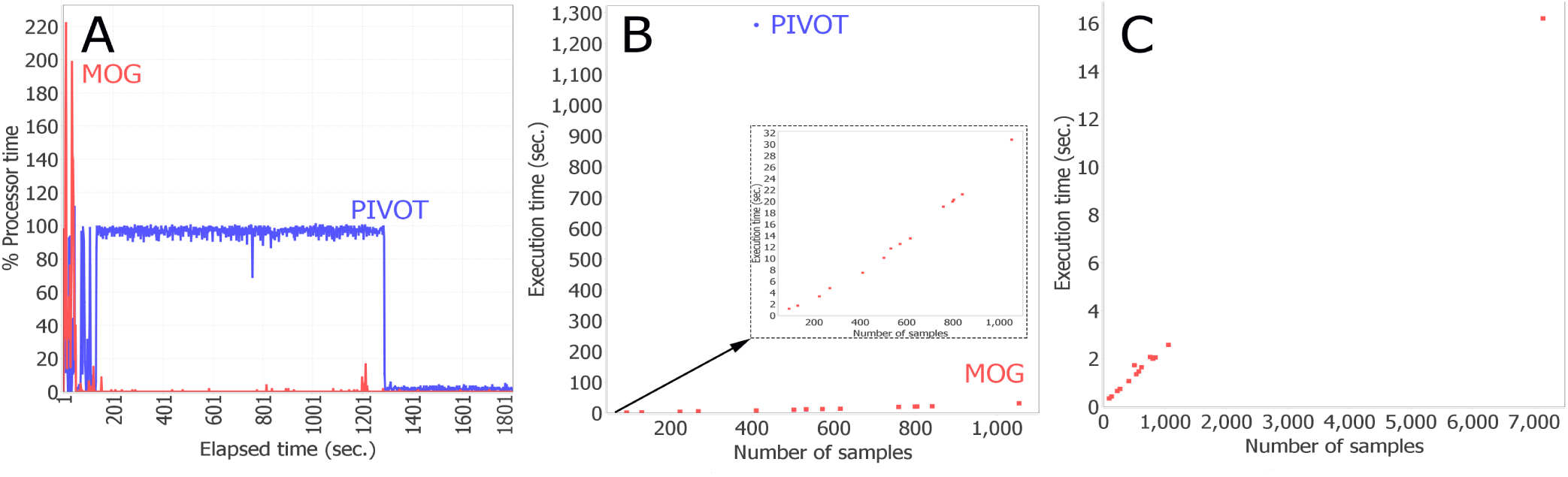
MOG performance benchmarks. MOG was benchmarked using the entire the cancer dataset (18,212 genes over 7142 samples) and using smaller-sized datasets derived by partitioning the full cancer dataset into tumor and non-tumor samples from various tissues.**(A)** Comparison of MOG to R-based (PIVOT). Dataset size was limited to the amount of data that could be loaded in PIVOT i.e., liver tumor and non-tumor samples (18,212 genes over 410 samples). The % processor time (% CPU utilization) for PIVOT and MOG was calculated over 30 minutes. The theoretical maximum value of % processor time is total processorsavailable in the computer × 100 (400 in this case). **(B)** Execution times for computing differentially-expressed genes using Mann-Whitney test with datasets of different sizes. Red dots, execution times for MOG; blue dot, execution time for PIVOT’s with 410 samples (as in panel **A**). Inset, same graph with expanded scale of Y-axis to display MOG execution times. **(C)** MOG execution times for pairwise computations of Pearson correlation of a gene (BIRC5) with all other genes in the datasets. (The other tools cannot perform this computation). Execution times are linear with data size, and analysis of the full dataset took just over 16 seconds.

### M. *Arabidopsis thaliana* metabolomics data

Metabolomics is providing a growing resource for understanding metabolic pathways and identifying the structural and regulatory genes that shape these pathways and their interconnected lattice (82, 126, 127). Here, we use a metabolomics dataset that represents a comprehensive study of 50 mutants with a normal morphological phenotype but altered metabolite levels, and 19 wild type control lines (81). There are 8-16 biological replicates for each genetic line; data is corrected for batch effects. Data and metadata were retrieved from PMR (82). The aim of this case study was to tease out coexpressed metabolites that are affected by genetic perturbations. We identified a group of four highly coexpressed metabolites (Pearson correlation *>* 0.8): the amino acid arginine, its precursor L-ornithine, cyclic ornithine (3-amino-piperidine-2-one), and one unidentified metabolite. Plots across the means of the biological replicates of each sample (Supplementary Figure 10), shows accumulation of these metabolites is upregulated over four-fold in four mutant lines: *mur9*, mutants have altered cell wall constituents; *vtc1*, encodes GDP-mannose pyrophosphorylase, required for synthesis of manose, major constituent of cell walls, upregulated upon bacterial infection; *cim13*, gene of unknown function associated with disease resistance, *eto1*, negative regulator of biosynthesis of the plant hormone ethylene. An arginine-derived metabolite, nitrous oxide, has been widely implicated in signaling pathways in plants (128). MOG analysis might suggest to a researcher a potential relationship between arginine and the cell wall defense response to consider for experimentation.

### N. Comparison to other software

Few tools that do not require coding are available for on-the-fly exploration of expression data. Most are essentially interfaces to R packages (18–20, 129), providing a “shiny” (17) web-app interface to a limited number of R packages for data normalization, differential expression analysis, PCA analysis (among samples) and gene enrichment analysis. Because they are written in R (18–20), these tools must rely on R’s somewhat limited capabilities for interactive applications. In contrast to R, Java, MOG’s platform, has been used to develop numerous software with interfaces that are interactive and userfriendly (102, 130–133), and MOG provides the researcher with specialized GUIs and methods for exploratory data analysis. MOG’s GUI allows direct interactivity with the data and the visualizations, so that a researcher can easily explore data from different perspectives. Table 4 compares four of the most recent tools for exploratory analysis of expression data to MOG. (More details are provided in Supplementary Table S34.)

**Table 4.** Comparison of MOG with existing tools for exploratory analysis of expression data. MOG’s GUI, designed with Java swing, is fully interactive. In contrast, other available tools are based on R and provide limited or no interactivity. With MOG, the user can execute any R package/script with interactively selected subsets of data if s/he wishes to perform additional analysis, whereas only a limited number of R-packages are available in the other packages. The last row compares the Mann-Whitney U test’s execution time for MOG and PIVOT using the liver tumor and non-tumor datasets (18,212 genes over 410 samples). A more detailed comparison of the tools is available in Supplementary Table S36.

|  | MOG | PIVOT | iGEAK | IRIS-EDA | DEvis |
| --- | --- | --- | --- | --- | --- |
| Reference | This paper | (18) | (19) | (20) | (129) |
| Year | 2019 | 2018 | 2019 | 2019 | 2019 |
| Platform/GUI | Java/Swing | R/Shiny | R/Shiny | R/Shiny | R/None |
| Interactivity | ++++ | + | + | + | - |
| Save progress | Yes | Yes | No | No | No |
| Use any R package | Yes | No | No | No | No |
| Supported data types | Omics or other numerical data | RNA-Seq/scRNA-Seq | RNA-Seq/microarray | RNA-Seq/scRNA-Seq | RNA-Seq |
| M-W U test (sec) | 7 | 1,260 | NA | NA | NA |

#### N.1. Benchmarking

We used the cancer dataset (18,212 genes over 7,142 samples) for benchmarking. Using a DELL Inspiron laptop with 64 bit Windows 10, 8 GB RAM and Intel(R) Core(TM) i5-7300HQ CPU, we monitored the system’s resource utilization using Windows Performance Monitor tool (WPMT) (134). During benchmarking, only the software being tested was run on the system and no other user program was running.

We compared MOGs efficiency to that of one of the “shiny” web-app (17) R software (choosing PIVOT, because it permits loading processed data). R-based tools read all data directly into the main memory. Thus, on a desktop computer, analysis of a big dataset is slow (or crashes) if the available memory is not sufficiently large. For example, a dataset of 100,000 human transcripts over 5,000 samples (500,000,000 expression values) requires at least 4GB (8 byte for each value) of free memory to be loaded into memory at once. In contrast, MOG uses an indexing strategy to read data only when it is needed, which drastically reduces the total memory consumption of the system.

PIVOT repeatedly crashed and failed to load the full (18,212 genes over 7,142 samples) cancer dataset: RStudio reported the error message: *“cannot allocate vector of size 992.4 Mb”* (Additional File 6). We then reduced the test dataset to only the liver tumor and non-tumor samples (18,212 genes over 410 samples), and were able to load this dataset to PIVOT. As a benchmark comparison of MOG and PIVOT, we performed a Mann-Whitney test to identify differentially-expressed genes in liver tumor and non-tumor samples and measured the execution time of this step, i.e., the time taken to produce and display the output. A Mann-Whitney test completed in 21 minutes using PIVOT, whereas MOG took only 7 seconds (Figure 9). After the Mann-Whitney test completed, we kept the program running idle until total runtime reached 30 minutes and compared memory and processor usage of PIVOT and MOG over this 30 minute window, as reported by WPMT (Supplementary Table S35-S36). Average memory usage of PIVOT (1,869 MB) was about twice that of MOG (995 MB) (Supplementary Table S35-S36). The peak % processor time of PIVOT’s when computing the Mann-Whitney test was about half that for MOG, indicating more CPU utilization by MOG; however, the average % processor time over 30 minutes was 64% for PIVOT but only 2% for MOG Figure 9 A).

We benchmarked MOG’s performance on datasets of different sizes, created by splitting the cancer dataset by different tissue types (tumor and non-tumor samples). For each dataset, we performed Mann-Whitney test on all the genes for tumor and non-tumor groups and measured the execution time. MOG took only 31 seconds to compute a Mann-Whitney test on 18,212 genes over 1,054 samples (Figure 9 B; Additional File 6).

We also measured the execution time for calculating the Pearson correlation of a gene (BIRC5 was chosen at random) with all others in a given dataset. MOG took only a couple of seconds to compute a Pearson correlation over 1,000 samples and 16 seconds to compute a Pearson correlation over all 7,142 samples (Figure 9 C).

## DISCUSSION

An advantage of MOG is that large datasets that have been normalized and unwanted technical or biological effects reduced (batch-corrected) based on different assumptions can be compared quickly and interactively by statistical and visual approaches, such that biologists can gain insight as to which best reflect experimentally-established “ground truths”. This approach is complementary to the tacts bioinfor-maticists usually take (e.g., assessing GO term enrichment in gene clusters). For the present, we have intentionally avoided incorporating capabilities for data normalization and batch-correction into MOG for two reasons.

First, the selection of appropriate methods depends on the data distribution and the biological questions to be asked. Different types of data have different characteristics and thus require different normalization methods (37, 52, 135). Removing/minimizing batch-effects before evaluating the data also entails more assumptions and approaches. These statistical methods can easily be misapplied, especially when the data is from multiple heterogeneous studies, resulting in a misleading dataset. Much as if using R or MATLAB statistical software, a MOG user must consider these. Second, the methodologies are very unsettled ((37, 51, 53, 135)) with new approaches and variations being developed each year (for RNA-Seq raw reads there were over 10,000 journal articles in first half of 2019 in GoogleScholar, searched on “RNA-Seq normalization methods”).

We demonstrated MOG’s functionality and detailed its application by exploring three different datasets: a human cancer RNA-Seq dataset from non-tumor and tumor samples, an *A. thaliana* microarray dataset, and an *A. thaliana* metabolomics dataset. Exploring the large human dataset, we created a catalog of genes which are differentially expressed in different types of cancers, identifying only a handful of genes that are consistently upregulated or downregulated in every type of cancer. Using the sample and study metadata, we identified genes that showed regulation with cancer progression. Comparing our results with genes described in THPA, indicated that some of the genes we identified have been reported in literature to be potential prognostic biomarkers for different cancers, however, many others of the genes are *not marked as prognostic in THPA*. These genes present potential new biomarkers for disease progression. Because each tumor type has many variations, investigating such markers in individual tumors would provide critical information for personalized medicine, as well as aid cancer research (136). GPC3 is known (93, 95, 96), and also was identified by MOG from the data, as a marker gene for liver cancer. Gene-level resolution analysis revealed that the genes that are coexpressed with GPC3 gene change drastically among the individual tissues, and between tumor samples and non-tumor samples.

Using the *A. thaliana* microarray dataset, we explored expression patterns of genes with unknown functions including orphan genes, demonstrating a plant-restricted gene that was tightly coexpressed with a set of genes involved in in photosystem assembly, and identifying an Arabidopsis specific gene that is highly expressed in pollen development.

## CONCLUSION

MOG is a novel tool that enables interactive exploratory analysis of big omics datasets; and indeed any normalized dataset can be input to MOG. Visualizations produced by MOG are fully interactive, and can easily allow researchers to detect and mine interesting data points. The statistical methods implemented in MOG help to guide exploration of hidden patterns in a user-friendly manner. By integrating metadata, MOG affords an opportunity to extract new insights about the data based on information about each feature, or any of the diverse information entered by experimenters about the biology of each sample and the experimental protocols and parameters used to obtain its data.

## DATA AVAILABILITY

MOG is free and open source software published under the MIT License. MOG and all compiled datasets in this article are freely downloadable from http://metnetweb.gdcb.iastate.edu/MetNet_MetaOmGraph.htm. MOG’s source code and user guide is available at https://github.com/urmi-21/MetaOmGraph/.

## SUPPLEMENTARY DATA

Supplementary data are available at BioRxiv and https://github.com/urmi-21/MetaOmGraphMOG_SupplementaryData. Additional files are available at https://github.com/urmi-21/MetaOmGraph/MOG_SupplementaryData.

### ACKNOWLEDGEMENTS

We especially thank Nick Ransom for his key role in MOG’s early development. We are grateful to our collaborators, Kevin Bassler, Pramesh Singh and Ling Li, for their help and feedback. We much appreciate the efforts of Jing Li, Priyanka Bhandhary, Arun Seetharam, and the early-adopters who beta-tested MOG and provided valuable feedback.

