## Supplementary Figures for "MetaOmGraph: a workbench for interactive exploratory data analysis of large expression datasets"

4 Modules identified in coexpression networks made using 3,012 differentially expressed genes in LIHC vs normal liver samples and visualized using cytoscape [2]. (**A**) Shows the GPC3 containing module identified in the network from liver samples. GPC3 is directly connected to 21 genes (shown in yellow). (**B**) Shows the module found in the network inferred from LIHC samples. In this network GPC3 was absent, but this module shared 33 genes with that of from the liver normal network. Genes which were directly connected to GPC3 in first module are shown in yellow. . . . . . 4

5 GO terms enriched in the GPC3 containing module (Supplementary Figure 4 A) found in the coexpression network from the liver samples visualized using REVIGO [1]. . . . . . . . . . . . . . . . . . . . . . . . . . . . . . . . . . . . . . . . . . . . . . . . . 5

6 GO terms enriched in the second module (Supplementary Figure 4 B) found in the coexpression network from the LIHC samples visualized using REVIGO [1]. . . . . . . 6

7 Differential analysis of genes upregulated in pollen versus leaf- sample selection. Pollen samples were selected as in the inset. Leaf samples were selected by leaf in organs and plant ontology. . . . . . . . . . . . . . . . . . . . . . . . . . . . . . . . . . . . . . 7

8 Differential expression analysis followed by volcano plot visualization using MOG. Gene metadata, revealed upon hovering the mouse over a data point, shows At3G25760, a gene of Embryophyta (photosynthetic land plants), is about 50-fold more highly accumulated in leaves relative to pollen (Mann-Whitney U test; B-H corrected p-value$< 10^{-16}$. Among these is At3G25760, specific to Arabidopsis and encoding a protein of unknown function; it is about 50-fold upregulated. A Pearson's correlation of this gene versus all genes across all samples, followed by a GO enrichment test of all genes with a correlation greater than 0.70, indicates the most significantly enriched genes encode enzymes of biosynthesis of jasmonic acid, a hormone that orchestrates injury response in plants [3], fungal-responses, and wounding responses. . . . . . . . . . 8

9 Genes coexpressed with AT1G67860 (Spearman correlation $> 0.65$) are highly expressed in leaf samples as compared to pollen. . . . . . . . . . 9

10 Upregulation of correlated metabolites in Arabidopsis. There is a significant increase in accumulation of these metabolites in four methylation mutants. Each data point is means of 8-16 biological replicates. (*transcribed ORFs*) genes. Metabolomics data from [Fukushima A, Kusano M,Mejia RF,Iwasa M, Kobayashi I,Hayashi N,Watanabe-Takahashi A,Narisawa T, Tohge T,Hur M,Wurtele ES,Nikolau BJ,Saito K. 2014. Metabolomic Characterization of Knock-Out Mutants in Arabidopsis - Development of a Metabolite Profiling Database for Knock-Out Mutants in Arabidopsis (MeKO). Plant Physiol. doi:10.1104/pp.114.240986] downloaded from PMR (http://metnetweb.gdcb.iastate.edu/PMR/) . . . . . . . . . . 10

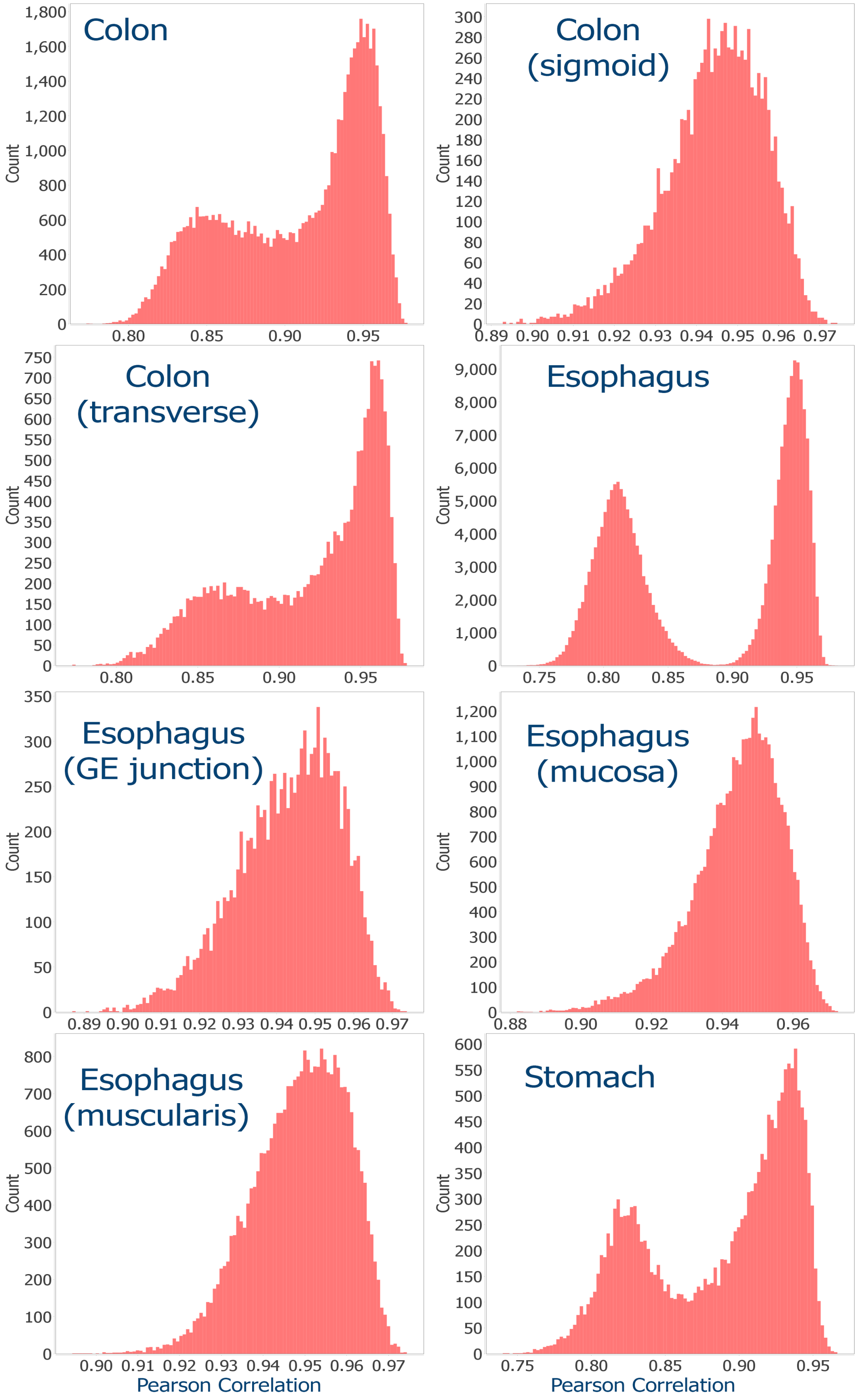

FIG. 1. The distribution of Pearson correlation values among samples from colon, esophagus and stomach tissues.

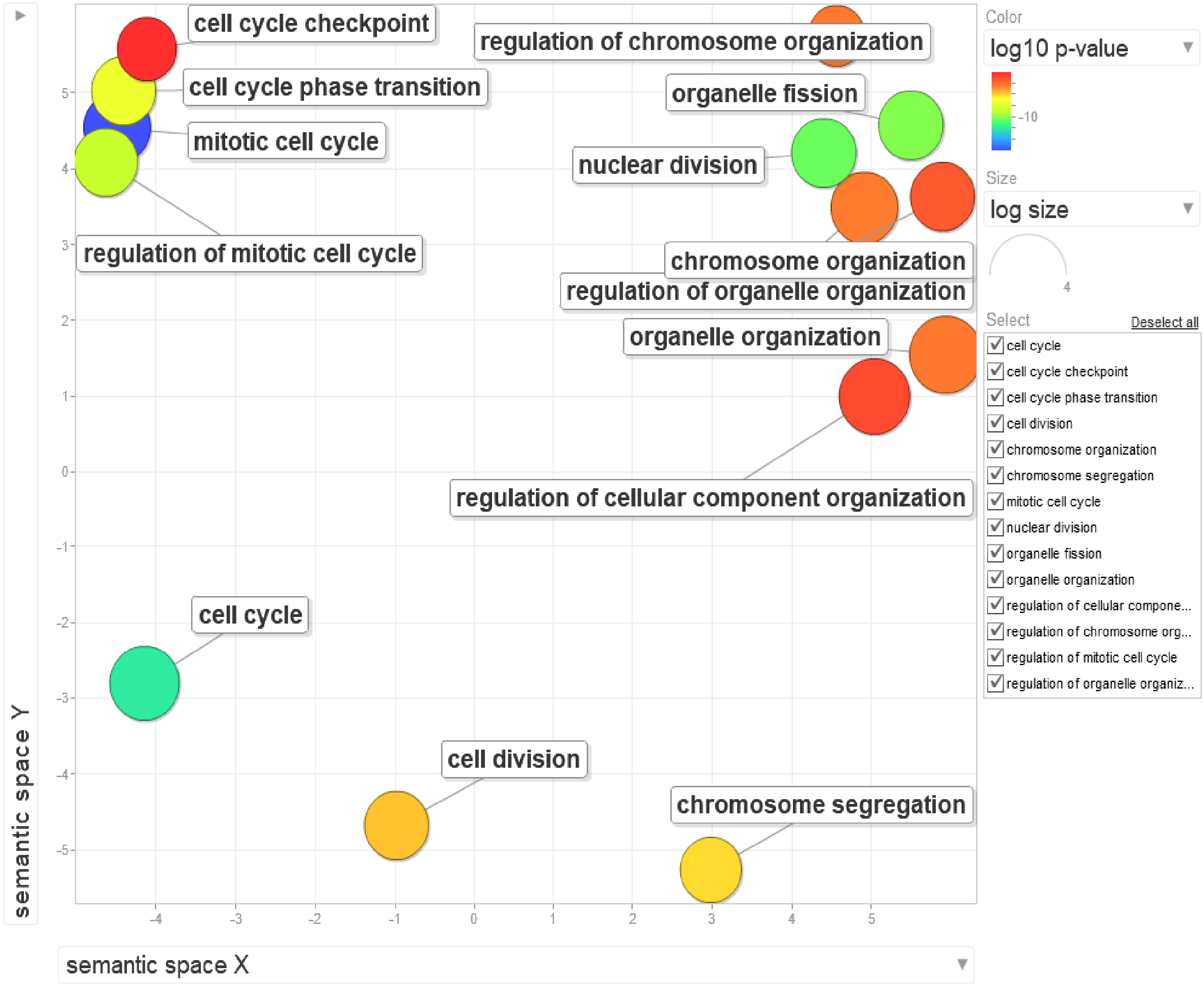

FIG. 2. GO terms enriched in the 16 upregulated genes in all 14 cancer types visualized using REVIGO [1].

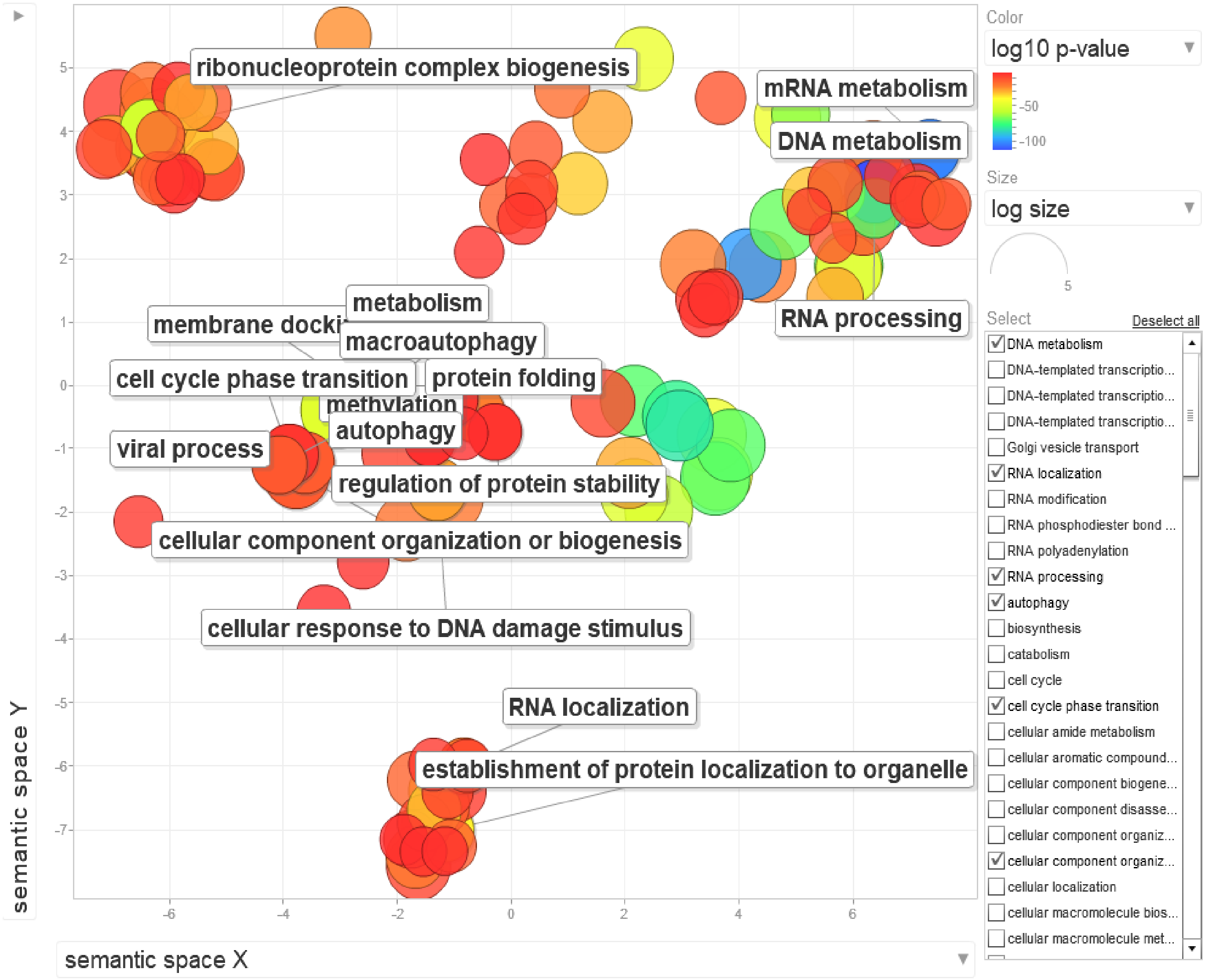

FIG. 3. GO terms enriched in the 5,784 unchanged genes in all 14 cancer types visualized using REVIGO [1].

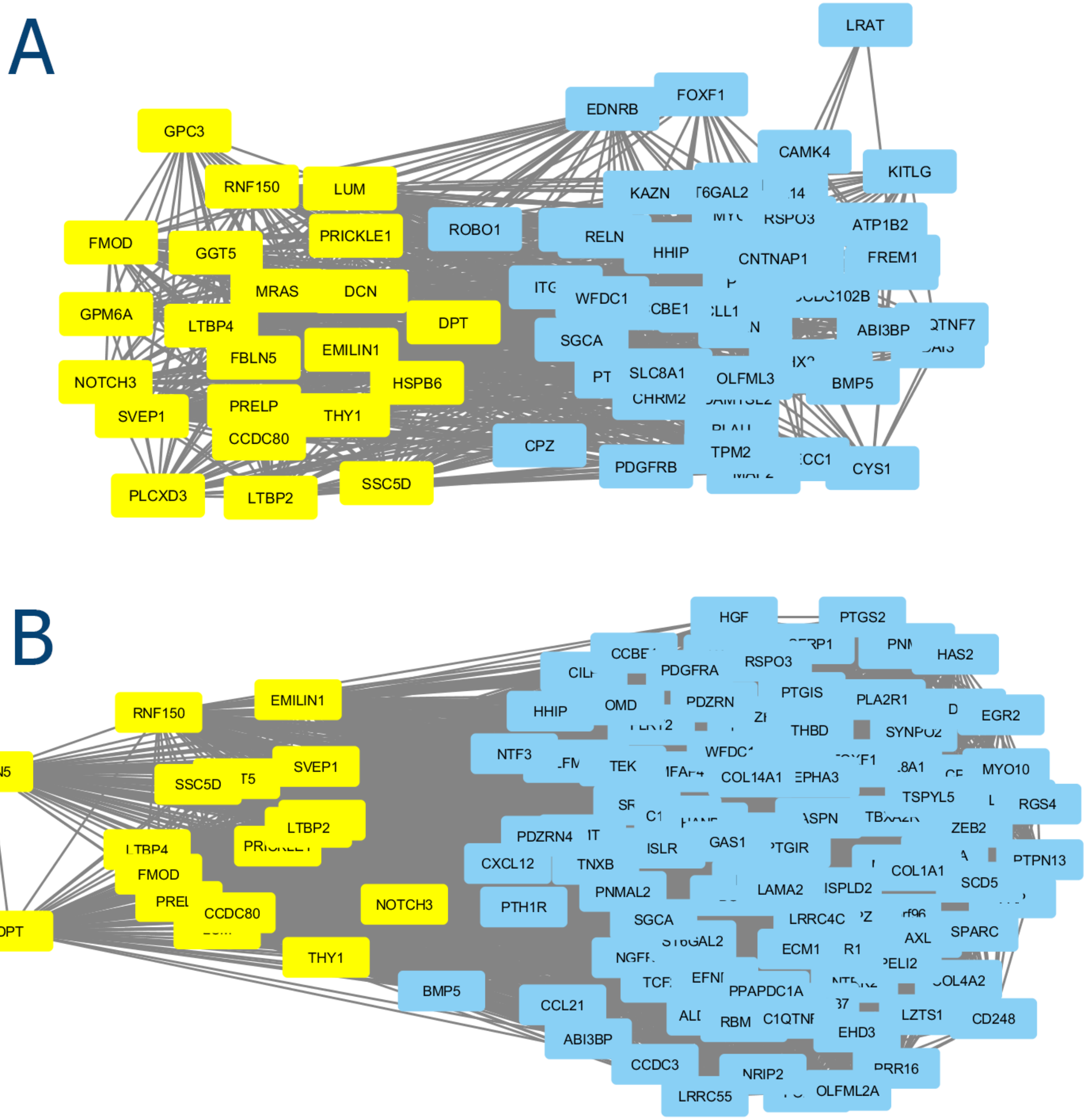

FIG. 4. Modules identified in coexpression networks made using 3,012 differentially expressed genes in LIHC vs normal liver samples and visualized using cytoscape [2]. (**A**) Shows the GPC3 containing module identified in the network from liver samples. GPC3 is directly connected to 21 genes (shown in yellow). (**B**) Shows the module found in the network inferred from LIHC samples. In this network GPC3 was absent, but this module shared 33 genes with that of from the liver normal network. Genes which were directly connected to GPC3 in first module are shown in yellow.

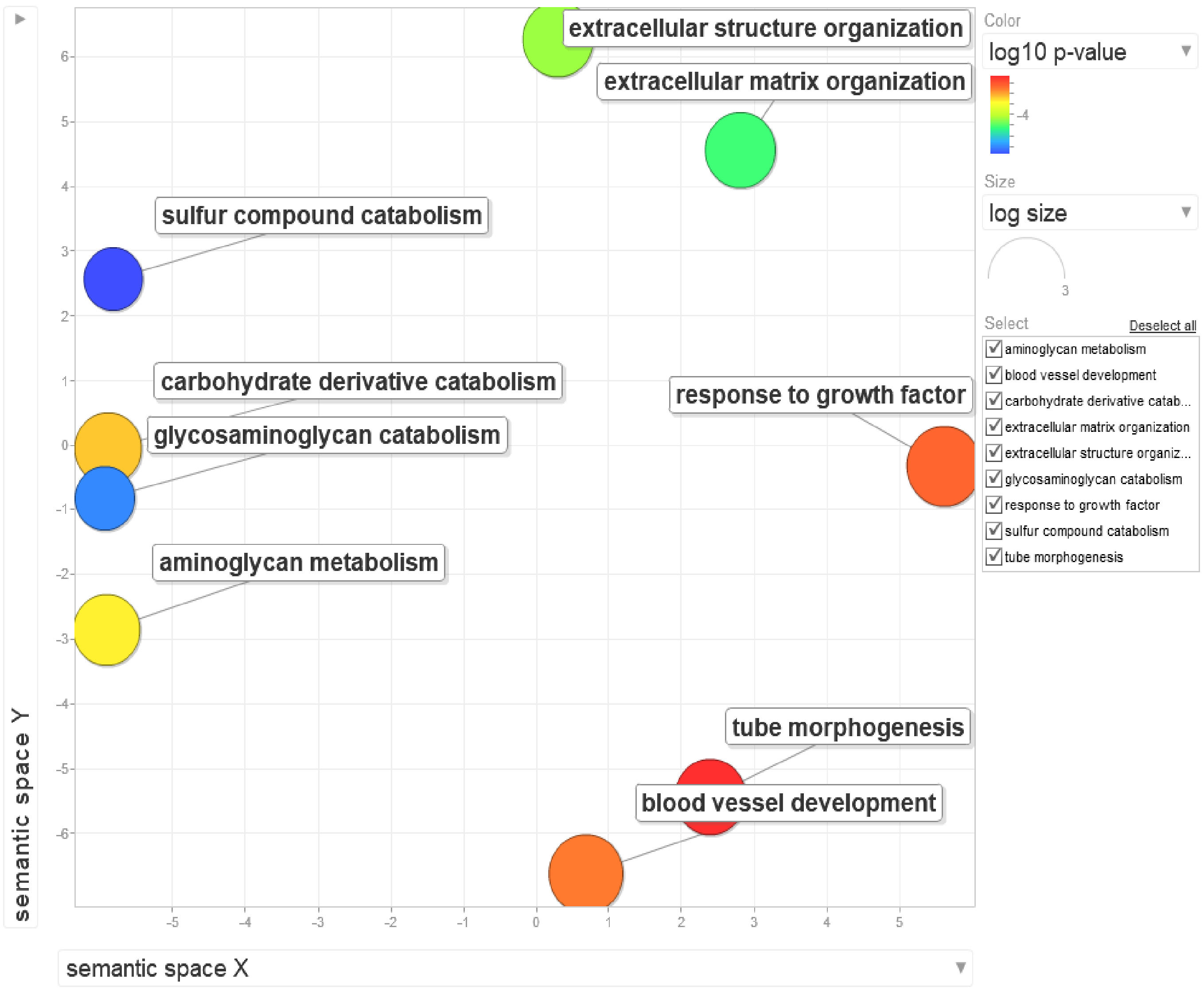

FIG. 5. GO terms enriched in the GPC3 containing module (Supplementary Figure 4 A) found in the coexpression network from the liver samples visualized using REVIGO [1].

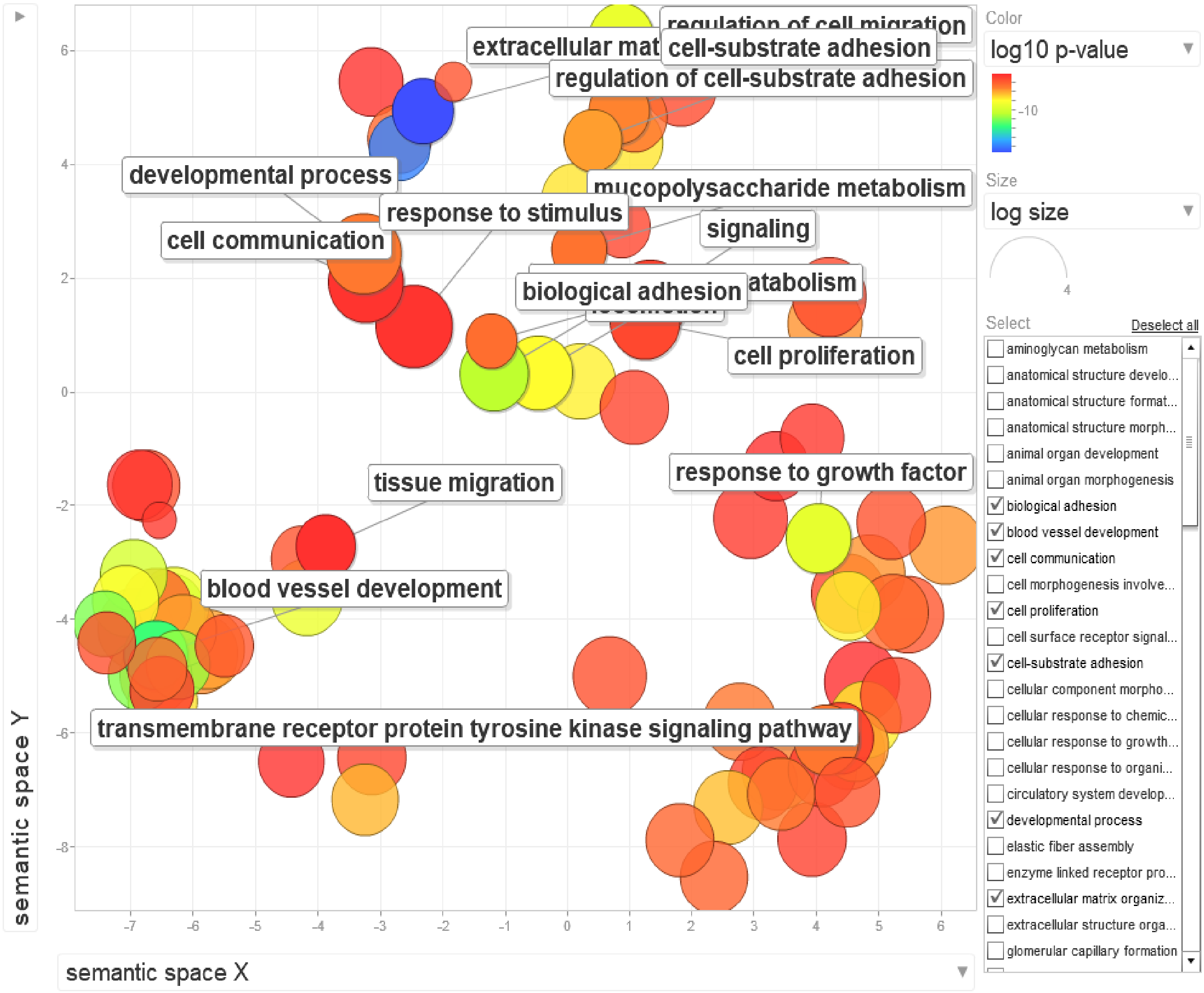

FIG. 6. GO terms enriched in the second module (Supplementary Figure 4 B) found in the coexpression network from the LIHC samples visualized using REVIGO [1].

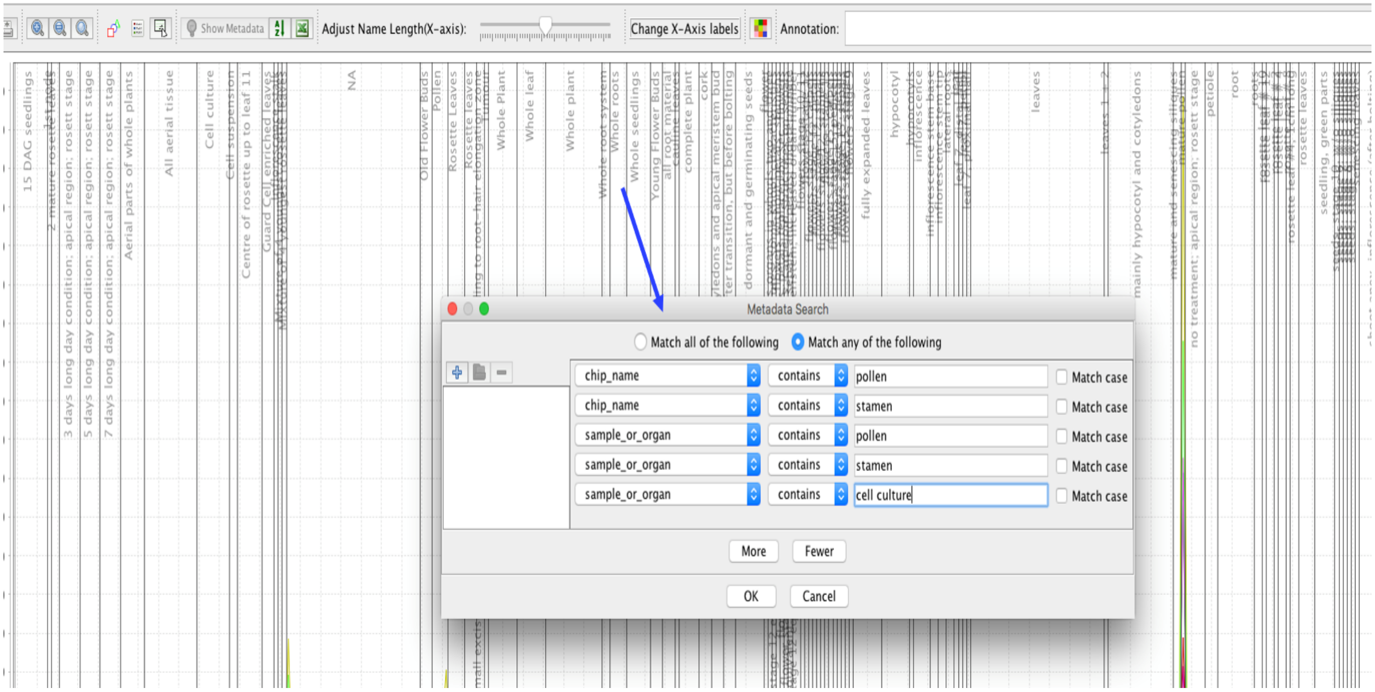

FIG. 7. Differential analysis of genes upregulated in pollen versus leaf- sample selection. Pollen samples were selected as in the inset. Leaf samples were selected by leaf in organs and plant ontology.

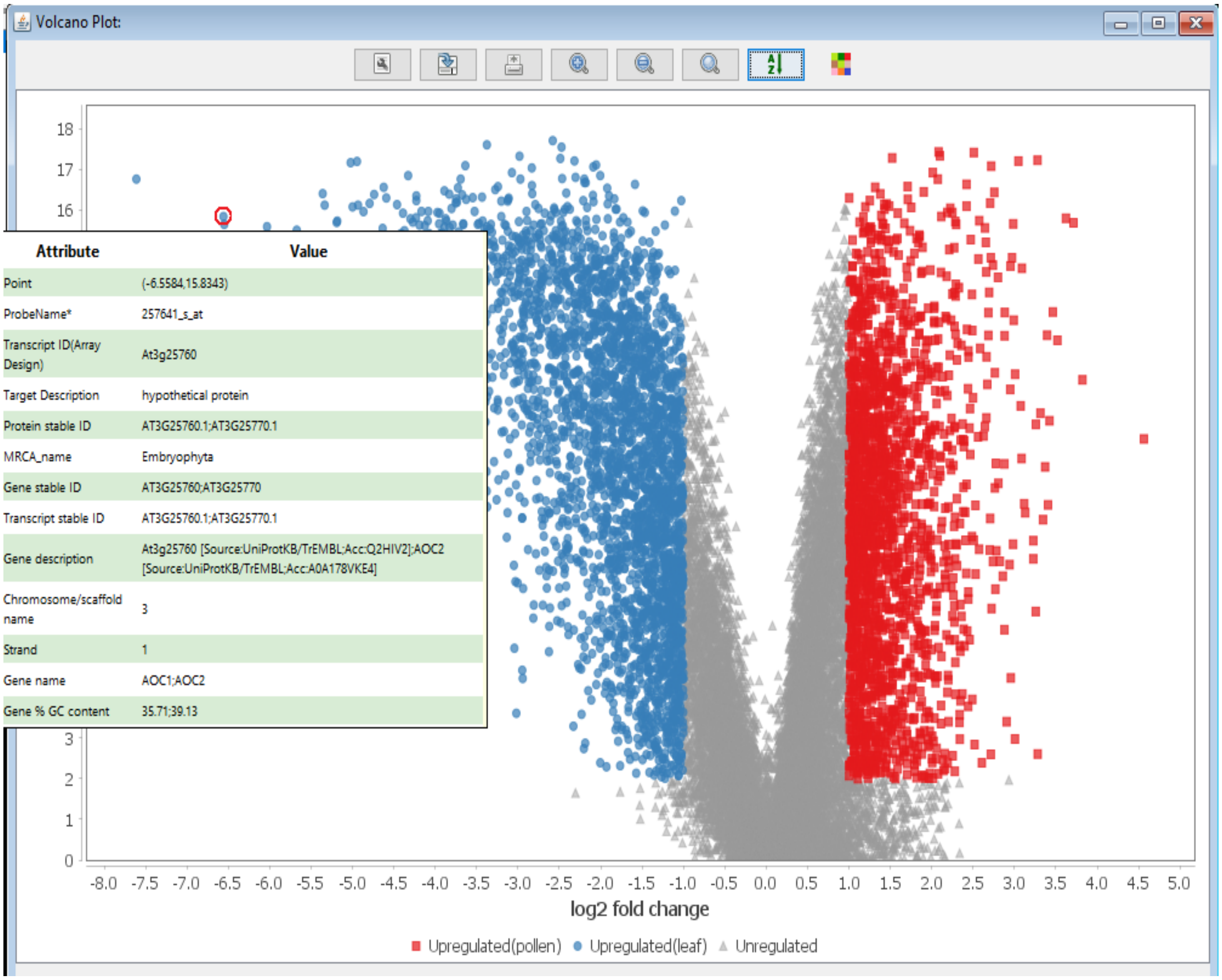

FIG. 8. Differential expression analysis followed by volcano plot visualization using MOG. Gene metadata, revealed upon hovering the mouse over a data point, shows At3G25760, a gene of Embryophyta (photosynthetic land plants), is about 50-fold more highly accumulated in leaves relative to pollen (Mann-Whitney U test; B-H corrected p-value$< 10^{-16}$. Among these is At3G25760, specific to Arabidopsis and encoding a protein of unknown function; it is about 50-fold upregulated. A Pearson's correlation of this gene versus all genes across all samples, followed by a GO enrichment test of all genes with a correlation greater than 0.70, indicates the most significantly enriched genes encode enzymes of biosynthesis of jasmonic acid, a hormone that orchestrates injury response in plants [3], fungal-responses, and wounding responses.

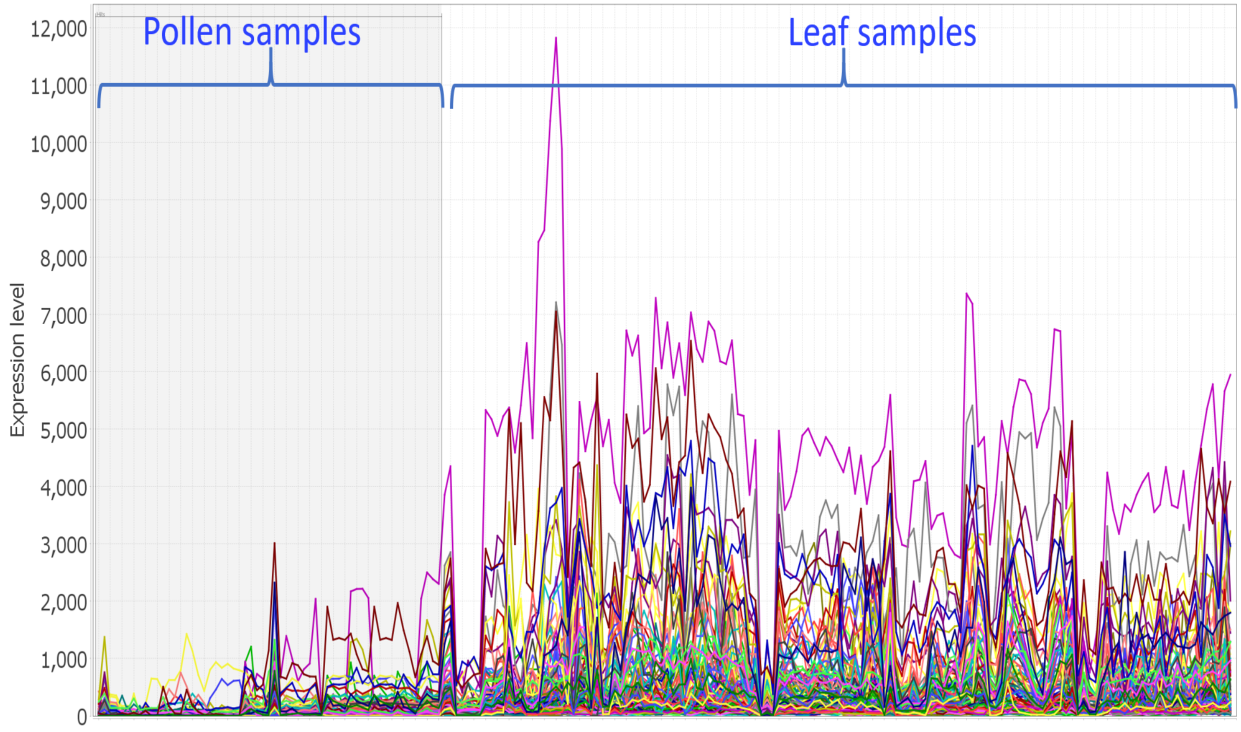

FIG. 9. Genes coexpressed with AT1G67860 (Spearman correlation > 0.65) are highly expressed in leaf samples as compared to pollen.

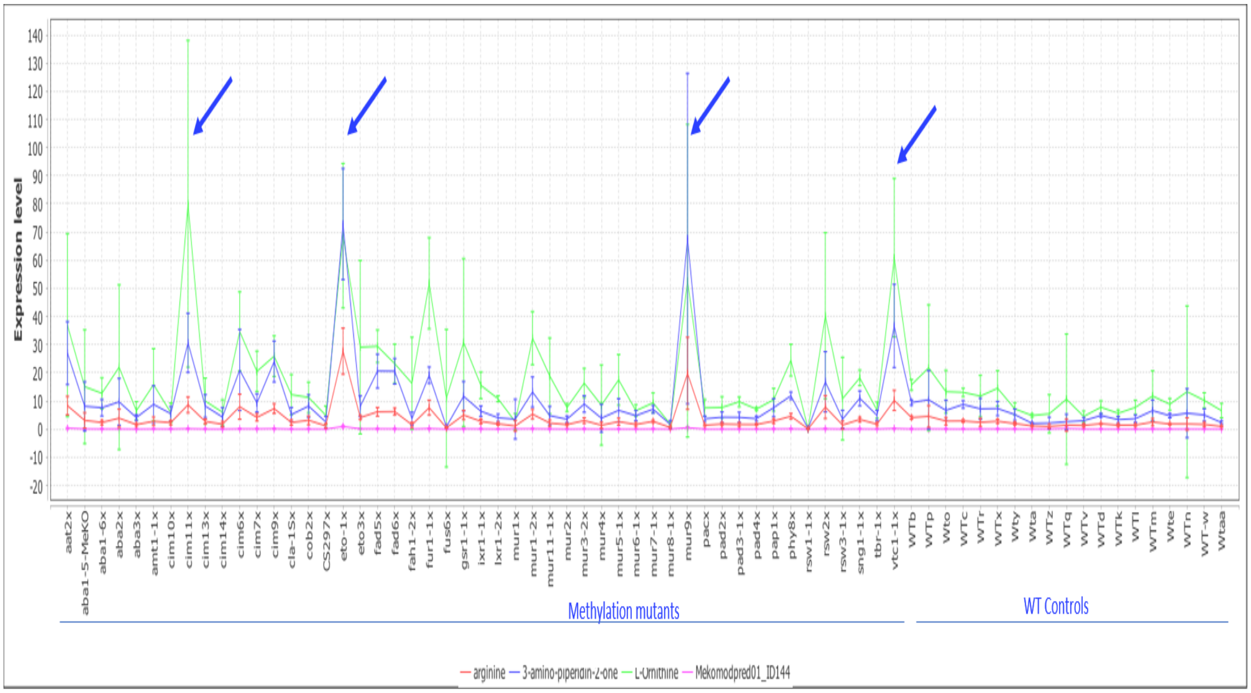

FIG. 10. Upregulation of correlated metabolites in Arabidopsis. There is a significant increase in accumulation of these metabolites in four methylation mutants. Each data point is means of 8-16 biological replicates. (*transcribed ORFs*) genes. Metabolomics data from [Fukushima A, Kusano M,Mejia RF,Iwasa M, Kobayashi I,Hayashi N,Watanabe-Takahashi A,Narisawa T, Tohge T,Hur M,Wurtele ES,Nikolau BJ,Saito K. 2014. Metabolomic Characterization of Knock-Out Mutants in Arabidopsis - Development of a Metabolite Profiling Database for Knock-Out Mutants in Arabidopsis (MeKO). Plant Physiol. doi:10.1104/pp.114.240986] downloaded from PMR (http://metnetweb.gdcb.iastate.edu/PMR/)

[1] Supek, F., Bošnjak, M., Škunca, N., and Šmuc, T. (2011) REVIGO summarizes and visualizes long lists of gene ontology terms. *PloS one,* **6**(7), e21800.

[2] Shannon, P., Markiel, A., Ozier, O., Baliga, N. S., Wang, J. T., Ramage, D., Amin, N., Schwikowski, B., and Ideker, T. (2003) Cytoscape: a software environment for integrated models of biomolecular interaction networks. *Genome research,* **13**(11), 2498–2504.

[3] Koo, A. J. and Howe, G. A. (2009) The wound hormone jasmonate. *Phytochemistry,* **70**(13-14), 1571–1580.
